# A New Paradigm of Transcriptional Regulation by the SufR-Like Iron-Sulfur Transcription Factors

**DOI:** 10.64898/2025.12.26.696629

**Authors:** Zhifang Lu, Chloe Ong, Lingwen Zhang, Tao Wan, Omar Davulcu, Marcelo de Farias, Limei Zhang

**Affiliations:** Department of Biochemistry, University of Nebraska-Lincoln, Lincoln, NE, USA; Pacific Northwest Center for Cryo-EM, Oregon Health & Science University, Portland, Oregon, USA; Pacific Northwest National Laboratory, Environmental Molecular Sciences Laboratory, Richland, Washington, USA; Redox Biology Center, University of Nebraska-Lincoln, Lincoln, NE, USA; Biortus Biosciences Co., Ltd, Jiangyin, Jiangsu, 214437, China

**Keywords:** SufR, iron–sulfur cluster, transcription factor, AT-hook, winged helix-turn-helix (wHTH), COG2345, ArnR

## Abstract

SufR is an iron–sulfur ([4Fe-4S]) cluster-containing transcription factor belonging to an uncharacterized domain family (COG2345). It has been shown to negatively regulate the sulfur utilization factor (SUF) Fe-S biogenesis system in several Gram-negative and Gram-positive bacteria including Cyanobacteria, Mycobacteria and Streptomyces. The structural basis for its DNA recognition and transcriptional regulation by the SufR-like proteins remains enigmatic. In this study, we present the cryo-EM structure of *Mycobacterium tuberculosis* SufR bound to its promoter, revealing a new domain architecture. Our structural, biochemical and molecular analyses show that SufR possesses an unusual [4Fe-4S] cluster coordination environment in the sensory domain and recognizes its promoter via a dual-module mechanism using both the AT-hook and the helix-turn-helix (HTH) DNA-binding motif in the DNA-binding domain. This DNA recognition strategy differs from a canonical winged HTH transcription factor. Moreover, our bioinformatic analysis and structural modeling suggest that SufR and SufR-like proteins in the COG2345 family represent a large, previously uncharacterized family of transcription factors widely distributed across bacteria and archaea. Together, these findings establish SufR-like proteins as a new model for transcriptional regulation by Fe-S transcription factors in prokaryotes and uncover the underappreciated evolutionary versatility of AT-hooks.

## Introduction

Iron-sulfur (Fe-S) clusters are ancient, versatile cofactors composed of multiple iron and sulfur ions. The tunable reactivity of Fe-S clusters makes them excellent candidates for sensing and rapidly responding to diverse environmental stimuli in bacteria (reviewed in ref. ^1,2^). To date, only a few Fe-S regulators have been characterized in details, including the fumarate and nitrate reduction regulatory protein FNR, the iron-sulfur cluster (ISC) biogenesis regulator IscR, the nitric oxide sensor NsrR, the redox-responsive regulator SoxR, and members in the WhiB-like family transcription factors (such as WhiB2 and WhiB7) ^1,3–10^. However, the diverse structures and mechanisms of transcriptional regulation by many other Fe-S transcription factors remain poorly understood, which includes the SufR-like transcription factors.

SufR is a [4Fe-4S] transcription factor belonging to the COG2345 family (annotated as “ArsR-like family transcription regulators”, Architecture ID: 11457150) in the National Center for Biotechnology Information (NCBI) Conserved Domain Database (CDD) ^11^. It was first reported to regulate the *suf* operon in *Synechocystis sp.* strain PCC 6803 (*Ssp*), and subsequently in other cyanobacterial species as well as in Mycobacteria and Streptomyces ^12–19^. In its [4Fe-4S] cluster-bound (holo-) form, the SufR homodimer represses transcription of the *suf* operon and thus modulates the maturation of many Fe-S proteins ^13,15,16^. SufR has also been implicated in response to nitric oxide stress, an important immune signaling pathway, and contributes to the survival and pathogenesis of *Mycobacterium tuberculosis* (*Mtb*) ^18^. Molecular and biochemical analyses suggest that SufR possesses a winged helix-turn-helix (wHTH) DNA-binding motif in the N-terminus and binds to a [4Fe-4S] cluster in the C-terminus via three conserved Cys (corresponding to C179, C192, C220 in *Mtb* SufR) and an unidentified non-Cys ligand (Supplementary Fig. S1) ^15–18^. DNase I footprinting and electrophoretic mobility shift (EMSA) assays have mapped the SufR binding site to a palindromic sequence (CAAC-N6-GTTG) in *Ssp* and *Streptomyces avermitilis* (*Sav*), and an imperfect palindromic sequence (ACACT-N5-TGTGA) in *Mtb* ^15–17^. However, the structural basis for DNA recognition and transcriptional repression by SufR remains elusive due to the lack of 3D structural information for SufR or any other proteins in the COG2345 family.

Here we report the single-particle cryo-electron microscopy (cryo-EM) structure of *Mtb* SufR bound to the promoter of the *suf* operon. Our structural and biochemical analyses reveal that SufR adopts a new domain architecture unlike any experimentally determined protein structures in the Protein Data Bank. Moreover, our structural and biochemical analyses show that SufR coordinates the [4Fe-4S] cluster with an unusual glutamate ligand and employs a rare DNA recognition mechanism in prokaryotes using an AT-hook in the wing region of the wHTH motif. The AT-hook motif, first identified in the high-mobility group protein (HMGAs), is a small DNA-binding motif (<10 residues) defined by a central “Gly-Arg-Pro [GRP]” sequence and often flanked by positively charged residues ^20–23^. It preferentially binds to the minor groove of A/T-rich DNA, commonly found in eukaryotes but considered rare in bacteria. Our bioinformatic analysis and structural modeling, however, show that SufR and SufR-like transcription factors in the COG2345 family are widespread in bacteria and archaea. Together, these findings unravel a new regulatory mechanism by the SufR-like Fe-S transcription factors in prokaryotes and uncover the previously unrecognized evolutionary versatility of the AT-hook motif.

## Results

### A/T-rich sequences in the promoter of the *suf* operon are required for SufR binding

Previous studies suggested that SufR binds to a palindromic sequence in the promoter of the *suf* operon (*P_sufR_*) in Cyanobacteria, Streptomyces and Mycobacteria ^15–17^. We noted that, in all cases, the identified SufR binding site is flanked by an A/T-rich sequence on both sides, as exemplified by *Mtb P_sufR_* (Fig. 1A). These A/T-rich sequences, consisting of consecutive adenine-thymine base pairs ([A/T]_n_, n≥4) and often referred to as A-tract DNAs, are well-known for their distinct structural features such as narrow minor grooves and high propeller twist angles comparing to the canonical B-form DNA ^24–27^. Such A/T-rich DNAs are recognizable by the AT-hook DNA-binding motif and play an important role in controlling gene expression, as demonstrated in previous studies on the antibiotic-responsive Fe-S transcription factor WhiB7 in *Mtb* ^8,9^. Consistently, sequence analysis reveals two Arg-rich motifs in the N-terminus of SufR homologs, including a central “GRP” AT-hook motif (Fig. 1B and C; Supplementary Fig. S1) ^13,16,17^. Actinobacterial SufR homologs share an invariant “RGRP” motif similar to WhiB7, whereas a non-polar residue (such as Met, Leu and Pro) often precedes the “GRP” motif in cyanobacterial homologs.

**Figure 1.**
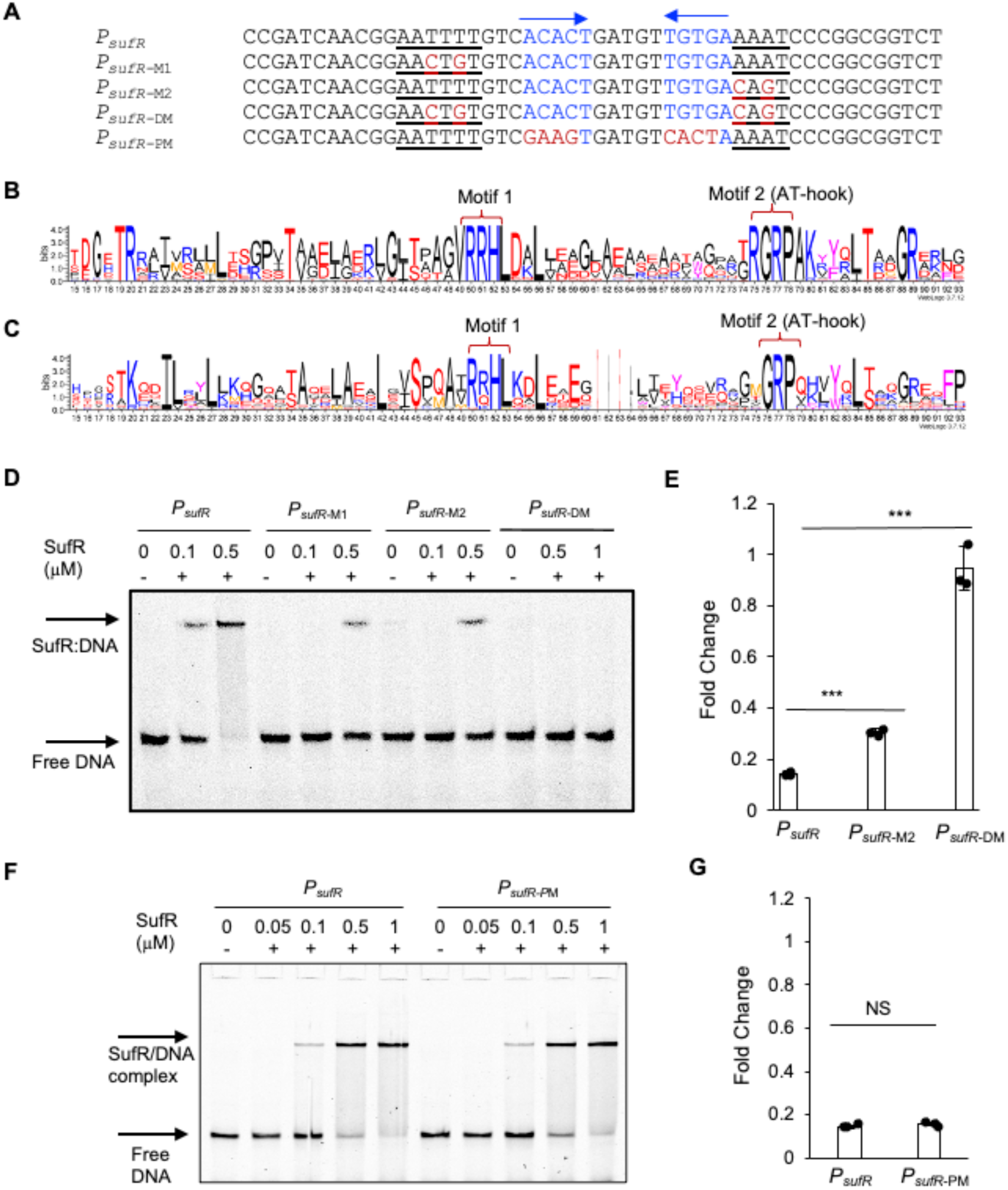
Characterization of the SufR binding site in the promoter of the *suf* operon. (A) The native and mutated promoter sequences of the *suf* operon (*P_sufR_)* are indicated. The previously identified palindromic sequence of the SufR binding site is marked in blue with arrows. The flanking A/T-rich regions are underlined. The mutated bases are highlighted in red. Panels (B) and (C) are the sequence logos of the N-terminal SufR homologs in Actinobacteria and Cyanobacteria, respectively. The full sequence logo is shown in Supplementary Fig. S1. Panels (D) and (F) are the electrophoretic mobility shift assays (EMSAs) of SufR binding to the native and mutated *P_sufR_* as shown in (A). Each reaction contained 1 nM Cy5-labeled *P_sufR_* DNA (wildtype and mutant, as indicated) and holo-SufR at the indicated concentrations. Panels (E) and (G) are the bioluminescence assays in *E. coli* using a heterologous lux-reporter under the control of either the wildtype or mutant *P_sufR_* as indicated (see Methods). The plasmid expressing the wildtype SufR was used in the assays, and the empty vector was used as the negative control. The fold change was calculated as the ratio of the bioluminescence measured in each testing sample relative to the negative control. Statistical significance was determined using Student’s *t* test (***, *P* < 0.001; NS, not significant) from three biological replicates.

We then carried out mutagenic analysis to test whether the A/T-rich region in the *P_sufR_* promoter is required for transcriptional regulation by SufR. As shown in Fig. 1D, disruption of the A/T-rich sequence on either side (*P_sufR_*-M1 and *P_sufR_*-M2, respectively) significantly reduces SufR binding to DNA in EMSAs. Consistently, the repression activity of SufR from the mutated *P_sufR_*-M2 is reduced by ∼25% compared to the wildtype promoter in the bioluminescence assay (see Methods, Fig. 1E) ^17,28^. Disruption of both flanking A/T-rich sequences completely abolishes DNA binding and transcriptional repression. Moreover, shortening the *P_sufR_* DNA from 50 bp to 31 bp (including only the palindromic sequence and the flanking A/T-rich sequences), as used in the previous report ^18^, leads to a significant decrease in binding affinity in the EMSAs, indicating the importance of the A/T-rich DNA structure for SufR binding (Supplementary Fig. S2A). In contrast, mutating the nucleotides in the core palindromic region of *P_sufR_* shows little effect on either DNA binding or transcriptional repression by SufR (Fig. 1F and G). These observations support that SufR exhibits low sequence specificity in the palindromic region and utilizes the flanking A/T-rich DNA sequence for promoter recognition.

We further quantified the binding affinity of *Mtb* SufR to the *sufR* promoter by MicroScale Thermophoresis (MST) using the same 50-bp *P_sufR_* DNA as was used in the EMSAs (see Methods, Fig. 1A, Supplementary Table S1) ^29^. As shown in Supplementary Fig. S2B, the MST assay shows a single binding event between SufR and *P_sufR_* with a dissociation constant (*K_d_*) of 234.9 ± 45.8 nM.

### Cryo-EM analysis reveals a new domain architecture in the DNA-bound SufR

To gain insight into the structural basis for how SufR recognizes its target DNA and represses transcription, we prepared the *Mtb* SufR:*P_sufR_* complex sample for cryo-EM analysis. The *Mtb* SufR:*P_sufR_* complex contains a 224-residue (residues 6-229) *Mtb* SufR dimer bound to a 50-bp *P_sufR_* DNA (see in Materials and Methods, Supplementary Table S1). The grids were prepared under a low O_2_ environment to protect the [4Fe-4S] cluster in SufR from oxidative degradation. We obtained a 3D reconstruction map at 3.5-Å average resolution (Supplementary Figs. S3 and S4). The initial SufR structural model was built using the Phenix PredictAndBuild tool ^30^, which integrates AlphaFold-based structural prediction and map-guided model construction, followed by structural refinement with the addition of the *P_sufR_* DNA and the [4Fe-4S] clusters in the model (see in Materials and Methods). The final structural model includes 222 residues of *Mtb* SufR and a 37-bp DNA fragment (see Supplementary Table S2 for statistics).

The cryo-EM structure reveals that each protomer in the SufR dimer consists of three domains: an N-terminal DNA-binding domain (DBD), a central dimerization domain (DD) and a C-terminal sensory domain (SD) containing the [4Fe-4S] cluster (Fig. 2A-D). They are related to each other by a C2-axis perpendicular to the DNA helix axis with a Cα root mean square deviation [RMSD_Cα_] of 0.68 Å over 212 residues. The two DBDs in the DNA-bound SufR adopt a domain-swapped configuration. The axis of inertia of the C-terminal domain is oriented at approximately 30° to the DNA helix axis (Fig. 2C).

**Figure 2.**
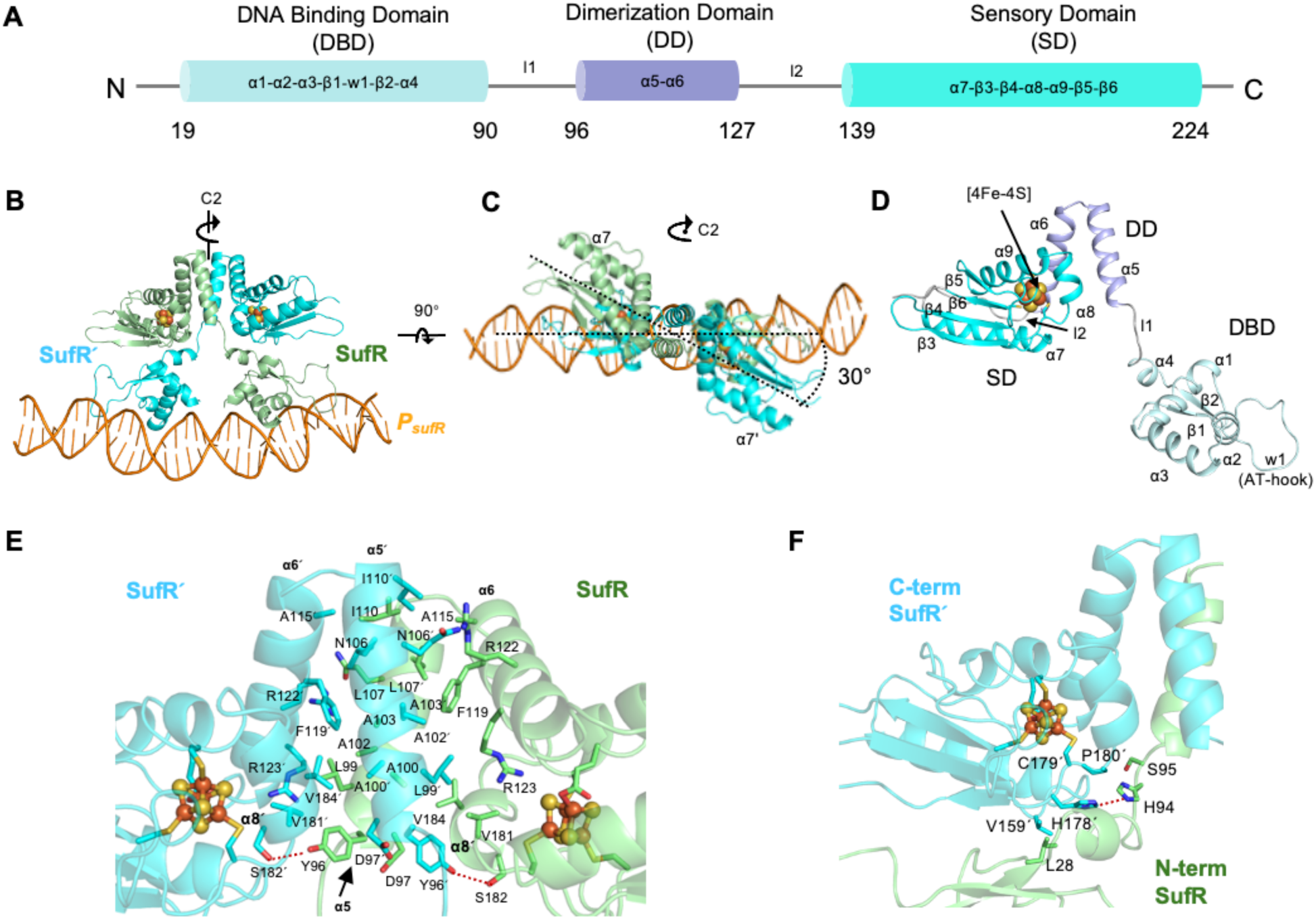
Overview of the SufR:*P_sufR_* cryo-EM structure. (A) Schematic diagram of the domain organization in *Mtb* SufR. (B) and (C) are cartoon representations of the SufR:*P_sufR_* complex. The two protomers in the SufR dimer (noted as SufR and SufŔ) are shown in palegreen and cyan, respectively, and the *P_sufR_* DNA is shown in orange. The pseudo-C2 axis for the SufR dimer is shown in the figure. (D) Cartoon representation of the domain organization in the SufR protomer, with the N-terminal DNA-binding domain (DBD) colored palecyan, dimerization domain (DD) in light blue and the C-terminal sensory domain (SD) in cyan. Panels (E) and (F) are the interactions between the dimerization domains and between the N- and C-termini of SufR and SufŔ, respectively. The cryo-EM density map for figures in (E) and (F) are shown in Supplementary Fig. S5. In all cases, the [4Fe-4S] clusters are highlighted in ball and sticks, and the residues at the dimerization interface are shown in sticks. The Fe atoms are colored red, S in orange, N in blue and O in red.

The SufR protomers (denoted as SufR and SufŔ) share a buried surface area (BSA) of ∼1638 Å^2^, with the primary dimerization interface between the two DDs (α5-6 and α5’-6’) and stabilized by extensive hydrophobic interactions between predominantly short-chained residues (Fig. 2E and Supplementary Fig. S5A). In addition to this DD–DD’ interface, helix α5 in DD also forms contacts with helix α8’ (residues 181’-184’) in the SD’, including a hydrogen bond between S182’ and Y96 (3.4 Å apart) (Fig. 2E). Y96 is highly conserved, and a polar residue is dominant at the S182’ position. The interactions between the DBD and the domain-swapped SD’ involve a combination of hydrophobic (L28···V159’), hydrophilic (H94···H178’) and van der Waals interactions (Fig. 2F and Supplementary Fig. S5B). Of note, H178 is adjacent to the Fe-S cluster-bound C179 and is highly conserved in the SufR homologs (Supplementary Fig. S1). H94 is not conserved, but a polar residue is dominant at the position. No contact is observed between the two DBDs in the cryo-EM structure of SufR:*P_sufR_*. The relative positioning of the DBDs relative to the other two domains is likely correlated with DNA-binding, given that each DBD is connected the rest of the protein by a flexible, disordered loop (*l1*) and does not make extensive domain–domain contacts (Fig. 2D and F).

The overall structural arrangement of the SufR DBD resembles that of a canonical wHTH domain found in many DNA-binding proteins, except that the winged loop (w1) is substantially longer and contains an AT-hook (Supplementary Fig. S5C). The DD and SD shares similarities in structural arrangement with a few known proteins despite of the low sequence identity, which include the sensory domain of the regulatory subunit of the trafficking protein particle complex subunit 3 (TRAPPC3, PDB ID:1WC8, 16% sequence identity) and the NtrC family transcription factor MopR (PDB IDs: 5KBG, 15% sequence identity) (Supplementary Fig. S5D) ^31,32^. However, a 3D structural search using FoldSeek shows no structural match of SufR in the Protein Data Bank that spans all three domains ^33,34^, reflecting the uniqueness of the domain-swapped DBDs in SufR.

### SufR uses a dual-module DNA recognition mechanism for transcriptional regulation

The local cryo-EM density map of the SufR:*P_sufR_* complex is well-resolved around the protein:DNA interface, allowing unambiguous assignment of the nucleotides and the DNA-binding residues (Figure 3A-D, Supplementary Fig. S6A-D). The structure reveals that the two conserved Arg-rich motifs in the SufR DBD, the “RRH” Motif 1 in helix α3 and the AT-hook in the w1 loop of the wHTH domain, play crucial roles in DNA binding (Figs. 1A-C and 3A). The AT-hook (residues 75-RGRP-78) slides into the minor groove of the A/T-rich region and makes extensive polar and nonpolar contacts with DNA, as previously observed in WhiB7 ^8,9^. In contrast, the helix α3 grips on the edge of the adjacent major groove and interacts primarily with the phosphate and sugar groups via the conserved “RRH” motif and S45. R50 and R51 also form van der Waals contacts with pyrimidines (T10 and C11’ for R50; T21 and T22’ for R51) in the major groove (Fig. 3D, Supplementary Fig. S6E-F). However, we did not observe any significant changes in binding affinity and repression activity of SufR when substituting C11’ with an adenine (Fig. 1F-G). Additional conserved residues in helix α1 (T19 and R20) of the wHTH domain also form contacts with the backbone of the major-groove DNA, likely enhancing the binding affinity. The mode of SufR binding to DNA strikingly differs from a typical wHTH transcription factor, in which the DNA recognition helix α3 inserts into the major groove to directly read out the nucleotide sequences (reviewed in ^35^). The results from the DNA structural analysis suggest that SufR binding induces conformational changes in *P_SufR_*. The central axis of *P_SufR_* bends ∼11° towards the SufR dimer (Supplementary Fig. S7A). Moreover, the central minor-groove width of *P_SufR_* around the AT-rich regions (∼6 Å) is significantly larger than that expected for A-tract DNA (typically smaller than 4 Å) and is consistent with previous reports of AT-hook-bound DNA (Supplementary Fig. S7B) ^9,20,21,36–38^.

**Figure 3.**
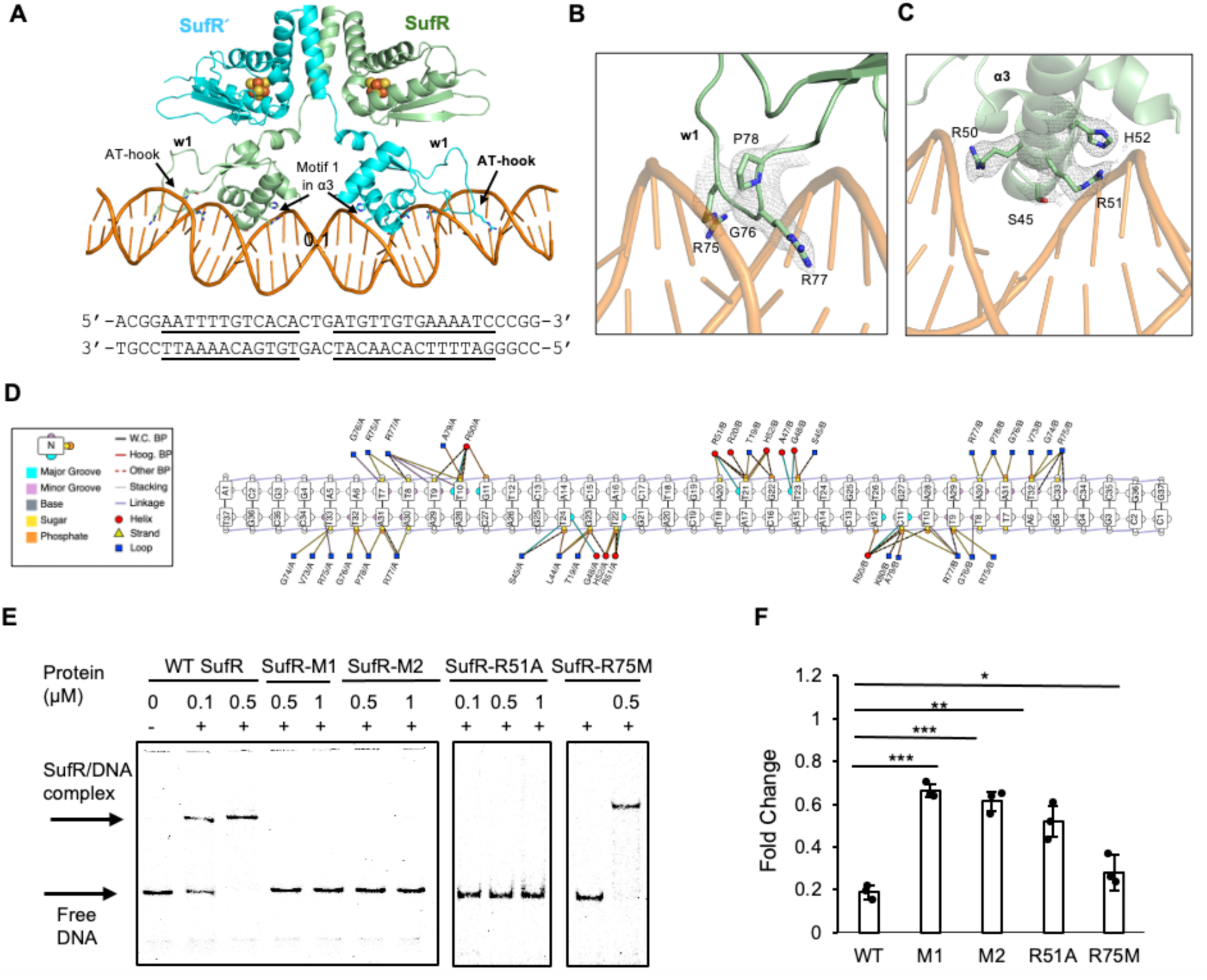
Interactions between SufR and DNA. (A) Overview of the SufR DNA-binding sites. The 37-bp DNA sequence in the SufR:*P_sufR_* complex cryo-EM structure is shown below the complex structure, with SufR’s binding sites underlined. (B) and (C) are the close-up views of the DNA-binding AT-hook in the w1 loop region and Motif 1 in helix α3 of the wHTH domain, respectively. The local cryo-EM density map around the DNA-binding residues is shown in gray and contoured at 2 σ. (D) A schematic presentation of DNA contacts by the AT-hook and the residues in the wHTH domain of SufR. The SufR-DNA contacts were analyzed using the online tool DBDproDB with a distance cutoff of 3.9 Å. The nucleotides and the contacting protein residues at the SufR:DNA interface are highlighted in color using the scheme shown in the legend and described in detail in the Methods. Hydrophilic interactions are shown as dashed lines, and the hydrophobic and Van der Waals interactions are shown as solid lines. (E) EMSAs of SufR (wildtype and mutant as indicated) binding to the 50-bp *P_sufR_* DNA. The uncropped EMSA images are shown in Supplementary Fig. S8. WT: wildtype; M1: the SufR mutant with the triple-alanine substitution of the “RRH” Motif 1 (50-RRH-50 -> 50-AAA-52); M2: the SufR mutant with the double-alanine substitution of the AT-hook motif (75-RGRP-78 -> 75-AGAP-78); R51A: the SufR mutant with the alanine substitution of R51 in the “RRH” motif; R51A: the SufR mutant with the methionine substitution of R75 mimicking the AT-hook motif in *Synechocystis* SufR. (F) Bioluminescence assays of the transcriptional repression from wildtype *P_sufR_* promoter by SufR (wildtype and mutant as indicated) in *E. coli* (see Methods). The same abbreviations for the SufR mutants are as described in Panel E. Statistical significance was determined using Student’s *t*-test (***, *P* < 0.001; **, *P* < 0.01; *, *P* < 0.05) from three biological replicates.

To evaluate the significance of the “RRH” Motif 1 and the “RGRP” AT-hook to DNA binding, we made alanine substitutions of the polar residues in each motif. As shown in Fig. 3E and F, either the triple-alanine substitution of the “RRH” motif (SufR-M1) or the double-alanine substitution of the Arg residues in the “RGRP” motif (SufR-M2) abolishes SufR binding to DNA *in vitro* under the experimental conditions and dramatically reduces its repression activity (42% and 48%, respectively, of the wildtype) in the bioluminescence reporter assays. An alanine substitution of R51, which forms extensive interactions with DNA, shows a similar effect on DNA binding and repression activity (59% of the wildtype), underlying its importance in the functional SufR. By contrast, substitution of R75 adjacent to the central “GRP’ AT-hook with a methionine (as in the AT-hook of *Ssp* SufR) shows little effect on DNA binding and repression activity (89% of the wildtype), consistent with the high variation of this residue in the sequence analysis. Of note, our western blot analysis shows that the cellular protein levels of all the mutants are comparable to the wildtype SufR (Supplementary Fig. S9), ruling out the possibility of reduced protein stability by any of these mutations. Together, these findings demonstrate that both DNA-binding motifs in the DBDs contribute significantly to DNA binding and repression activity of SufR.

### SufR binds to the [4Fe-4S] cluster via an unusual glutamate ligand

The cryo-EM structure reveals that the [4Fe-4S] cluster in the DNA-bound SufR is buried in the SD and coordinated by a glutamate (E195) and three previously identified cysteines (C179, C192, C220), all of which are strictly conserved in SufR homologs (Fig. 4A-B, Supplementary Fig. S10A). The local environment of the Fe-S cluster binding pocket outside the first-coordination shell is dominated by short-chain hydrophobic residues surrounding the cysteine ligands (Fig. 4B). In contrast, E195 forms a hydrogen bond with Q176 and is close to Y158 and R209 (< 4 Å). Although none of these three residues is conserved, a polar residue (most commonly Glu) is preferred at the Q176 position among the SufR homologs.

**Figure 4.**
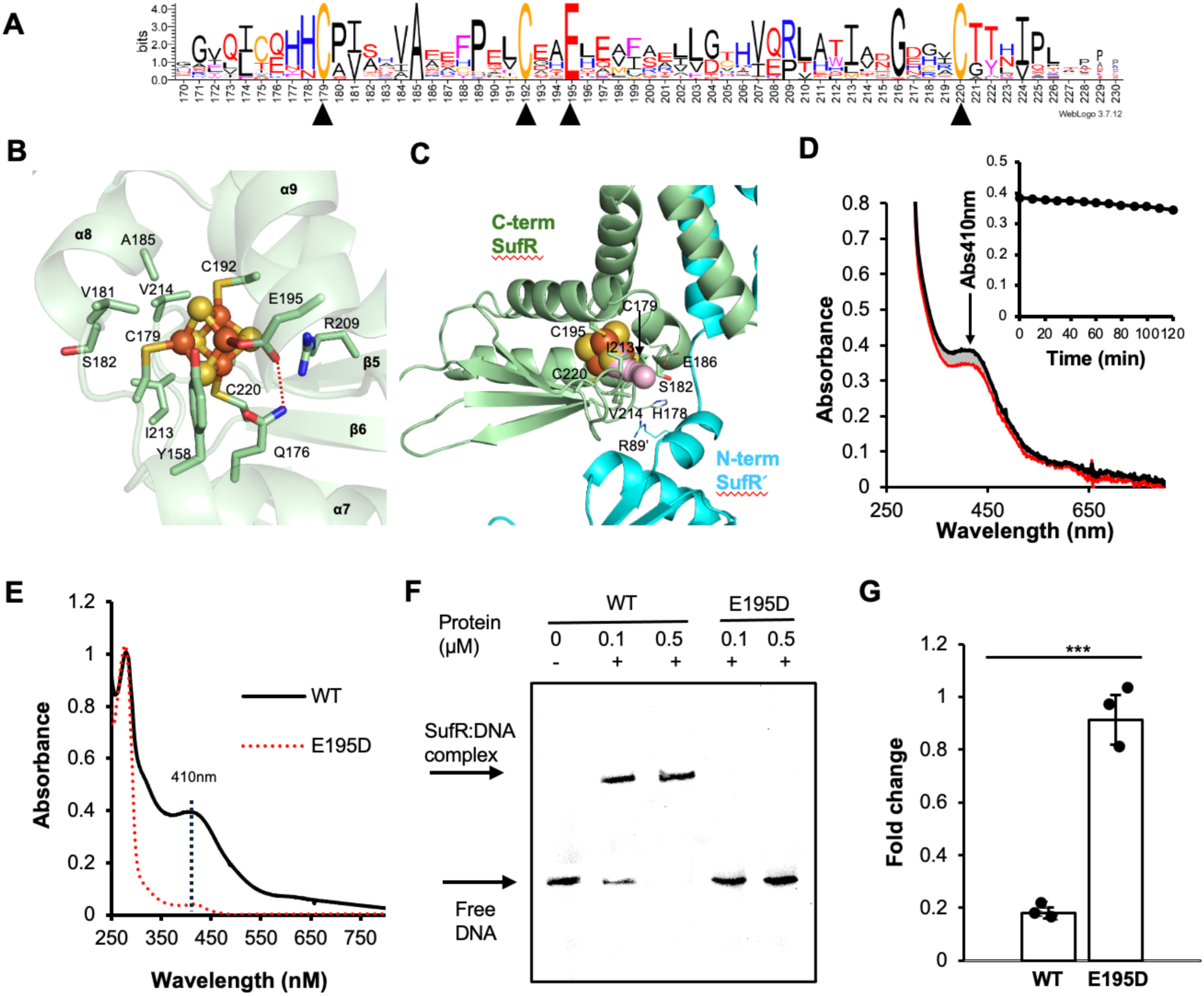
The local environment at the [4Fe-4S] cluster binding pocket of the *P_sufR_*-bound SufR. (A) Sequence logo of the C-terminal sensory domain of the SufR homologs. The [4Fe-4S] cluster-ligating residues in SufR are indicated by black arrowheads. The full sequence logo is shown in Supplementary Fig. S1. (B) Close-up view of the local coordination environment around the [4Fe-4S] cluster binding pocket. The [4Fe-4S] clusters are highlighted in ball and sticks. The cluster-binding residues and the residues surrounding the 1^st^-coordination shell of the cluster are highlighted in sticks. The averaged distance between E195-OE1 and Q176-NE2 is 3.08±0.13 Å, indicated by the red dashed line. (C) The tunnel from the solvent surface towards the [4Fe-4S] cluster in the SufR:*P_sufR_* complex calculated by CAVER using a probe radius of 1.0 Å. The minimum bottleneck of the tunnel is 1.16 Å. (D) Monitoring the [4Fe-4S] cluster loss in the DNA-bound SufR by UV-visible absorption spectroscopy upon exposure to O_2_ at 0 min (black curve), between 10-110 min (gray curve) and at 120 min (red curve), as indicated by a decrease in the absorption at 410 nm (highlighted by the black arrow). Inset, the plot of the absorption at 410 nm as a function of time. (E) Comparison of the UV-visible absorption spectra of wildtype SufR and the E195D mutant. The lower absorption intensity at 410 nm of the mutant is indicative of the cluster loss. (F) EMSAs of SufR (wildtype and E195A as indicated) binding to the 50-bp *P_sufR_* DNA. (G) Bioluminescence assays of the transcriptional repression from the wildtype *P_sufR_* promoter by SufR (wildtype [WT] and the E195A mutant as indicated) in *E. coli*. Statistical significance was determined using Student’s *t*-test (***, *P* < 0.001) from three biological replicates.

Cavity analysis indicates that the Fe-S cluster in the DNA-bound SufR is largely shielded from solvent, assuming a 1.4-Å radius cutoff for water molecules (Fig. 4C) ^39^. The largest tunnel from the solvent surface to the cluster is located at the interface between the DBD and SD’ of SufR and near Cys179, with a minimum bottleneck of 1.16 Å. Such limited solvent accessibility suggests that the [4Fe-4S] cluster in the DNA-bound SufR is resistant to O_2_ degradation, as previously reported ^40,41^. Consistently, we observe a very slow cluster degradation of the DNA-bound SufR under aerobic conditions, as indicated by minimal decrease of the absorption at 410 nm (∼5% in an hour) (Fig. 4D). By comparison, the free SufR loses its cluster ∼10 times faster under the same experimental conditions (Supplementary Fig. S10B), suggesting that DNA binding alters the conformation or dynamics in SufR and thus renders the O_2_ stability of the cluster.

E195 is located in helix α8, adjacent to the cluster-ligating residue C192, suggesting that the coordination geometry in this region is relatively rigid. Consistent with our structural analysis, a E195D mutation in SufR leads to a complete loss of the Fe-S cluster in the purified protein and abolishes its repression activity in the bioluminescence assay, even though the protein level of the mutant is comparable to the wildtype (Fig. 4E-G, Supplementary Fig. S9).

### SufR-like proteins represent a new class of transcription factors widespread in prokaryotes

SufR is classified in the COG2345 family, an uncharacterized domain family spanning multiple domains, in our NCBI Conserved Domain Database analysis. To assess the diversity and distribution of SufR and SufR-like proteins in prokaryotes, we analyzed the 326 representative members of the COG2345 family (see in Materials and Methods, Supplementary DataSet 1).

Results from sequence analysis and structural modeling reveal that the functional motifs and domain architecture of SufR are widely conserved among the representative COG2345 members despite the low sequence identity. In particular, the central “GRP” AT-hook motif in the DBD of SufR is highly conserved (Fig. 5A, Supplementary Table 3). R50 and H52 in the helix α3 of the wHTH domain of *Mtb* SufR are also conserved among most members, albeit with noticeable variations. Similarly, the three cysteines in the first coordination shell of the [4Fe-4S] cluster in the SD of SufR are found nearly all the representative COG2345 members with only five exceptions (Supplementary Table 3; Supplementary DataSet1). The cluster-ligating E195 in *Mtb* SufR is less conserved, but a residue capable of metal binding (such as Glu, His, Asn, Asp and Cys) predominates at this position (>96% of the sequences) (Fig. 5A, Supplementary Table 3). Together, these findings support that SufR-like proteins in the COG2345 family function as redox-responsive transcription factors and employ the dual-motif mechanism for DNA recognition similar to that of SufR, although we cannot definitely determine whether all of these members bind to a [4Fe-4S] cluster based on this analysis alone.

**Figure 5.**
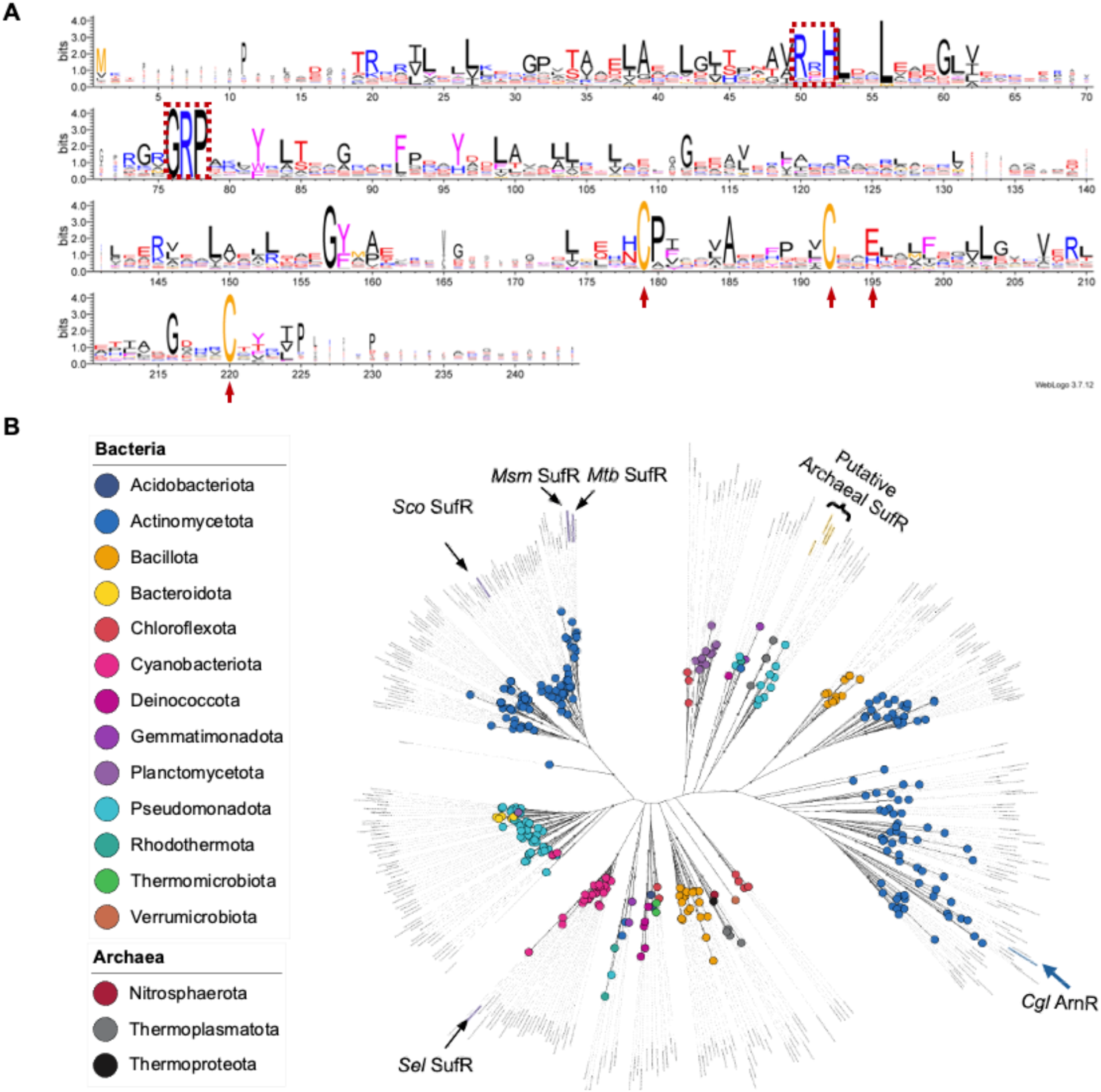
Sequence and phylogenetic analysis of representative members in the COG2345 family. (A) Sequence logo of the 326 representative COG2345 family sequences aligned against *Mtb* SufR. The numbering of the residues in the sequence logo follows *Mtb* SufR, with the alignment gaps in the sequence logos removed for clarity. The residues involved in DNA binding (the “50-RRH-52” motif and the central “75-GRP-78” AT-hook motif) and [4Fe-4S] cluster binding (C179, C192, E195 and C220) in *Mtb* SufR are highlighted by the red frame and arrows, respectively. (B) The unrooted phylogenetic tree of 326 representative sequences curated in the COG2345 family (Supplementary DataSet 1). The identified SufR proteins from Mycobacteria (*Mycobacterium tuberculosis* [*Mtb*] H37Rv and *Mycobacterium smegmatis* MC^2^ 155 [*Msm*]), Streptomyces (*Streptomyces coelicolor A3(2)* [*Sco*], encoding a close homolog of SufR from *Streptomyces avermitilis*) and Cyanobacteria (*Synechococcus elongatus* [*Sel*], encoding a close homolog of SufR from *Synechocystis sp.* PCC 6803) are highlighted purple on the tree. The putative archaeal SufR homologs highlighted in yellow on the tree were identified based on the conserved functional motifs and 3D structural models when compared to *Mtb* SufR and the location of the genes encoding these proteins immediately upstream of the *suf* operon. ArnR from *Corynebacterium glutamicum* (*Cgl*) is highlighted in blue.

Among the non-SufR COG2345 members in our analysis, only the aerobic repressor of nitrate reductase R (ArnR) from *Corynebacterium glutamicum* (*Cgl*) has known regulatory targets. *Cgl* ArnR shares 19.2% sequence identity with *Mtb* SufR and binds to a [4Fe-4S] cluster in its active form. Like SufR, ArnR functions as a transcriptional repressor, inhibiting the expression of the *narKGHJI* operon encoding enzymes involved in nitrate respiration, as well as the *hmp* gene encoding a flavohemoglobin involved in nitric oxide detoxification ^42–44^. Notably, the reported DNase I footprinting assays revealed that the protected region by ArnR on the *narK* promoter (*P_narK_*) DNA contains two A/T-rich sequences spanning ∼30 bp, despite the unexpected short consensus DNA binding site (10 bp) identified for ArnR (Supplementary Fig. 11A) ^42^. Consistently, the AlphaFold model of the *P_narK_*-bound ArnR closely resembles SufR with respect to the domain architecture and mode of DNA binding (see in Materials and Methods, Supplementary Fig. 11B). In particular, the AT-hooks in the ArnR dimer interact with the A/T-rich sequences in *P_narK_*, as observed in the cryo-EM structure of SufR:*P_sufR_*. The results from our EMSA assays further confirm the importance of the A/T-rich sequences in *P_narK_*-for ArnR binding (Supplementary Fig. 11C). As the rest COG2345 members are uncharacterized, we generated the AlphaFold models of these COG2345 members and compared them with SufR. It is worth noting that the AlphaFold models of the *P_sufR_*-bound *Mtb* SufR are essentially identical except for the N-term and C-term disordered region (Supplementary Fig. 12A), and they are closely resembles that of the cryo-EM structure of SufR:*P_sufR_* (RMSD_Cα_ = 1.14 Å over 212 aligned residues). In contrast, the AlphaFold models of *Mtb* SufR alone (as well as the other COG2345 members) vary significantly in the orientation of the DBDs relative to the other domains (Supplementary Fig. 12B), which reflects the highly dynamic nature of the loop region connecting the DBD and DD as described above in our structural analysis of SufR:*P_sufR_* and/or the uncertainty of the *ab initio* modeling of this region in the DNA-free form. Nonetheless, results from our 3D structural alignment suggest that these representative COG2345 members adapt a similar fold in the DBD and SD domains as SufR in the cryo-EM structure of SufR:*P_sufR_* and share an overall domain architecture comparable to that of SufR with only a few exceptions (Supplementary Fig. 13, Supplementary DataSet 1).

Phylogenetic analysis shows that the representative COG2345 family members are distributed across major bacterial phyla (Proteobacteria, Firmicutes, Actinobacteria, Cyanobacteria and Bacteroidetes) as well as in archaea (Fig. 5B). Besides the previously characterized SufR from Mycobacteria, Streptomyces and Cyanobacteria, we have also identified a cluster of uncharacterized archaeal proteins in the tree that contain the conserved functional motifs of SufR and are located immediately upstream of the *suf* operon (Fig. 5B), suggesting they are likely SufR homologs. Moreover, we note that many species in our analysis encode multiple COG2345 family members, *i.e.,* three each in *Mtb* (Actinobacteria), *Chroococcidiopsis thermalis* (Cyanobacteria) and *Cuniculiplasma divulgatum* (Archaea) (Supplementary DataSet 1), which share low sequence identity. These findings suggest that the COG2345 family members regulate diverse pathways beyond the SUF Fe-S cluster biogenesis.

## Discussion

SufR has been known as an [4Fe-4S] transcription factor to regulate the SUF Fe-S biogenesis system in bacteria since 2004. However, structural and mechanistic understanding of transcriptional regulation by SufR is lacking. In this work, we report the first atomic view of *Mtb* SufR bound to its promoter, revealing an unprecedented domain architecture and a new mechanism of transcriptional regulation by SufR and SufR-like Fe-S proteins that are widespread in prokaryotes and regulate different biological processes.

Importantly, this work expands our understanding of the role and distribution of the AT-hook in prokaryotic transcriptional regulation. The AT-hook is a short DNA-binding motif with relatively low binding affinity and specificity by itself. It is commonly found in eukaryotic DNA binding proteins, coordinating DNA recognition with another DNA-binding motif either within the same protein or the functional partner ^20–23^. By comparison, only two AT-hook proteins, WhiB7 and another transcription factor CarD from *Myxococcus xanthus,* have been identified in bacteria ^45–47^, and none in archaea thus far. Our analysis of SufR and SufR-like COG2345 family proteins provides compelling evidence that AT-hook proteins are far more common in prokaryotes than previously recognized and are involved in regulating diverse biological processes.

The results from our study support that SufR recognizes its target promoters primarily by specific DNA structural features rather than strict base sequence specificity and underscore the essentiality of the A/T-rich minor groove structure in DNA recognition by SufR. The spacing between the A/T-rich sequences also appears important, as a previous study showed that deletion of five nucleotides in this region abolishes SufR binding ^18^. These findings will be informative for identifying DNA binding sites of transcription factors with an AT-hook or AT-hook-like minor groove anchor. The conventional approach used for identifying the consensus DNA binding sites primarily focuses on nucleotide sequence specificity, while such an approach may overlook structure-dependent recognition elements as exemplified by SufR.

The glutamate (E195) at the [4Fe-4S] cluster binding site of SufR is a rare ligand of Fe-S clusters and has only been reported in two structurally characterized proteins ^48,49^: the hydrogenase-maturation enzyme HydF from *Thermosipho melanesiensis* (PDB ID: 5KH0), which coordinates a [4Fe-4S] cluster with three Cys and one labile glutamate; and the Rfr2-family transcription factor RrsR (PDB ID: 6HSD), which binds to a [2Fe-2S] cluster with two Cys, one His and one Glu. The cluster-ligating glutamate is highly conserved among the SufR homologs in Actinobacteria and Cyanobacteria in our sequence analysis (Fig. 4A), while it is replaced by a His in *Cgl* ArnR and its close homologs (Supplementary Fig. 11A) ^44^. The significance of this non-cysteine ligand on the Fe-S cluster’s reactivity and signal transduction of SufR and SufR-like proteins requires further investigation.

Another important question in the future study is how the [4Fe-4S] cluster regulates SufR’s transcriptional activity. Previous studies showed that loss of the cluster inactivates SufR without disrupting its dimeric structure ^15,18^, consistent with the strong hydrophobic interactions between the DDs of SufR revealed in our cryo-EM analysis (Fig. 2E). By contrast, the interactions between the DBD and the domain-swapped SD’, which involve the residues near the cluster-ligated C179 (Figs. 2F and 3), are relatively weak but could be crucial for stabilizing the DBD conformations and positioning them for DNA binding. The work reported here thus sets the foundation for further structural and mechanistic studies on [4Fe-4S] cluster-dependent DNA binding and transcriptional regulation by SufR and SufR-like transcription factors in prokaryotes.

## Materials and Methods

### Plasmids, oligos and bacterial strains

The bacterial strains, plasmids and oligos used in this study are listed in Supplementary Table S1. All the *E. coli* strains used for cloning and protein over-expression were grown in Luria–Bertani (LB) media supplemented with suitable antibiotics as needed at 37 °C with shaking at 200 rpm, unless otherwise specified.

All the oligos used in this study were synthesized by Millipore Sigma. An equal molar ratio of the paired oligos were annealed in annealing buffer (10 mM Tris, pH 7.5, 50 mM NaCl) and stored at -80 °C before use.

### Cloning for protein expression in *E. coli*

The open reading frame (ORF) of *Mtb sufR* gene has been experimentally examined and re-annotated to start at +73 bp located downstream of the start site annotated in Tuberculist (http://tuberculist.epfl.ch), resulting in a 240-aa protein, similar to the case of the *sufR* gene in *Synechocystis sp.* PCC 6803 ^15,50^. This revised ORF was used in this study. To improve the protein yield and stability, a DNA fragment encoding a truncated *Mtb* SufR including 6-229 aa (*Mtb* SufR_6-229_) was amplified from *Mtb* H37Rv genomic DNA, and subsequently cloned into the NcoI/XhoI site of pET28a (+) to express *Mtb* SufR_6-229_ with a C-terminal 6His-tag (*Mtb* SufR_6-229_-His_6_) for cryo-EM structural analysis and biophysical characterization. The resulting plasmid, pET28a SufR_6-229_-His_6,_ was further modified by site-directed mutagenesis to express mutant proteins as indicated. All the constructed plasmids were confirmed by DNA sequencing.

### Protein purification and analysis

The SufR proteins (wildtype and mutants) used for structural and biochemical studies were over-expressed in *E. coli BL21(DE3) Suf^++^*strain ^51^. The overexpression and anaerobic purification were carried out as previously described with minor modifications ^52^. Briefly, the cell pellets with overexpressed *Mtb* SufR_6-229_-His_6_ proteins (wildtype or mutant) were resuspended in lysis buffer containing 25 mM Tris pH 7.5, 150 mM NaCl, 50 mM arginine, 50 mM glutamate, 1 mM DTT, 10 µg/ml lysozyme, 3 µg/ml DNase I, and the cOmplete^™^ EDTA-free protease inhibitor cocktail (Millipore Sigma) per manufacturer’s instruction. After cell lysis by sonication and centrifugation, the whole-cell supernatant was loaded onto a Ni-NTA column (Cytiva Life Sciences). After washing with 5 column volumes (CV) of buffer containing 25 mM Tris pH 7.5, 1 M NaCl and 1 mM DTT and the 5 CV of buffer containing 25 mM Tris pH 7.5, 150 mM NaCl, 50 mM arginine, 50 mM glutamate, 30 mM imidazole and 1 mM DTT, the protein was eluted with buffer containing 25 mM Tris, pH 7.5, 150 mM NaCl, 50 mM arginine, 50 mM glutamate, 1 mM DTT and 300 mM imidazole. The eluted protein sample was further purified by passing through a Superdex 200 column (Cytiva Life Sciences) pre-equilibrated in the elution buffer containing 25 mM Tris, pH 7.5, 150 mM NaCl, 50 mM arginine, 50 mM glutamate and 1 mM DTT. The samples after each step of purification were analyzed by SDS-PAGE and by UV-visible spectroscopy. Unless otherwise specified, the final purified proteins in the elution buffer were stored in liquid nitrogen until use.

### Bioluminescence reporter assay in *E. coli*

The lux-reporter plasmid pOsufRlux, a kind gift from *Prof. Zhi Chen* (China Agricultural University), carries the *lux* operon (*luxCDABE*) that encodes a luciferase enzyme and the enzymes required for the biosynthesis of the luciferase substrate ^17,28,53^. This plasmid was modified to encode an ampicillin resistance gene, and a 248-bp promoter region of *Mtb suf* operon (genomic region 1,645,886-1,646,133, GenBank accession number: NC_000962.3) containing the native SufR binding site (*P_sufR_*) was cloned into the BamHI/XhoI site of the plasmid upstream of the *lux* operon to regulate its expression. To test the sequence specificity of the SufR binding site, the DNA sequence around the putative SufR binding site in the *suf* promoter region in pCS-*PsufR*-Lux-Amp was mutated to construct the variant reporters (*PsufR-*M2, *PsufR-*DM and *PsufR-*PM as listed in Supplementary Table S1). The resulting reporter plasmids, pCS-*PsufR*-Lux-Amp (either wildtype or mutant as indicated), were co-transformed into *E. coli* JM109(DE3) with either the empty vector pET28a (+) (negative control) or a SufR-expressing plasmid (pET28a-MtbSufR_6-229_-His_6_ or the mutated variants to express either wildtype or mutant SufR as indicated) to test the transcriptional repression activity of the SufR proteins.

For the bioluminescence assay, the fresh cultures of *E. coli* JM109(DE3) strains carrying a reporter plasmid and either the SufR expressing plasmid or the empty vector in late log phase (OD600nm of ∼1 – 1.5) were sub-cultured into a 96-well plate in LB media with the required antibiotics at OD_600nm_ of 0.15 and incubated at 37 °C, 200 rpm for 1.5 hr before data collection. The bioluminescence and cell density (OD_600nm_) were measured using a plate reader luminometer (CLARIOstar Plus, BMG Labtech). The bioluminescence readings were normalized by the cell density. All the assays were carried out with three biological replicates. Expression of the SufR proteins was from the leaky expression of the T7 polymerase without the addition of any inducer during the assay.

### Western Blot analysis

Cell cultures of *E. coli* JM109(DE3) strains expressing either the wildtype or mutant *Mtb* SufR_6-229_-His_6_ protein in the bioluminescence assays were harvested in late log phase (OD_600nm_ of ∼1 – 1.5) without induction. After cell lysis and centrifugation, the whole-cell supernatant samples containing 20 µg total protein each, as determined using the Pierce™ BCA protein assay kit (Thermo Scientific), were used for western blot analysis. The His-tagged SufR proteins were detected by chemiluminescence using mouse anti-His antibody (Thermo Scientific) as the primary antibody and horseradish peroxidase-coupled goat anti-mouse IgG antibody (Jackson Immuno Research Laboratories) as the secondary antibody in the western blot analysis.

### Electrophoretic mobility shift assays (EMSAs)

A Cy5-labeled 50-bp *sufR* promoter DNA (*P_sufR_*) (see the native and mutated *P_sufR_* sequence in Supplementary Table S1) was used in test SufR binding by EMSAs. For each assay, 1 nM DNA (final concentration) was incubated with varying concentrations of *Mtb* SufR_6-229_-His_6_ as indicated in the binding buffer (25 mM Tris-HCl, pH 7.5, 1 mM EDTA, 100 mM KCl, 100 mM NaCl, 1 mM DTT, 0.25 mg/ml BSA) under anaerobic conditions. 50 ng/µl Poly (dI•dC) was included in all reactions as a non-specific competitor. After 30-minute incubation on ice, 10 µl of the reaction mixture was loaded onto a 10% native polyacrylamide gel for electrophoretic separation. Gels were visualized using a Typhoon FLA 9500 biomolecular imager (GE Healthcare).

In the EMSA assays of Cgl ArnR and DNA, 50 nM of the 43-bp *narK* promoter DNA (*P_narK_*) (see the native and mutated *P_narK_* sequence in Supplementary Table S1) was used in the reaction, and carried out similarly as described above.

### Microscale thermophoresis (MST) analysis

MST measurements were conducted using a Monolith NT.115 green/red instrument (NanoTemper Technologies). A 50-bp *P_sufR_* DNA (*P_sufR_*) labeled with Cy5 at the 5’ end was used in the MST analysis (see the *P_sufR_* sequence in Supplementary Table S1). Cy5-labeled DNA (10 nM) was titrated with increasing concentrations of *Mtb* SufR_6-229_-His_6_ in binding buffer (20 mM Tris-HCl, pH 7.5, 100 mM KCl, 1 mM DTT) under anaerobic conditions in a COY anaerobic chamber (COY Laboratory). The final reaction mixture contained 5 nM Cy5-labeled DNA and the concentrations of SufR protein ranged from 100 µM to 3.05 nM. The reaction mixtures were incubated at room temperature for 30 minutes, centrifuged at 14k rpm for 2 min before loading into the standard capillaries (NanoTemper Technologies). All the capillaries were sealed with grease before being taken out of the anaerobic chamber for the MST measurements.

The MST measurements were done using the red channel LED source with 90% LED and 60% infrared-laser power at 25°C. The curve fitting and the *K_d_* value of the triplicate measurements were performed using MO Affinity Analysis v2.3 software (NanoTemper Technologies).

### Cryo-EM sample preparation, data collection and processing

To prepare the SufR:*P_sufR_* complex sample for cryo-EM, *Mtb* SufR_6-229_-His_6_ was incubated with a 50-bp *P_sufR_* DNA duplex (*P_sufR50_*, see the full sequence listed in Table S1) in a molar ratio of 1.5:1 in the buffer containing 25 mM Tris pH 7.5, 150 mM NaCl, 50 mM arginine, 50 mM glutamate and 1 mM DTT and then loaded onto a Superdex S75 10/300 column (Cytiva) to isolate the SufR:*P_sufR_* complex. The fractions containing the SufR:*P_sufR_* complex were combined and concentrated to ∼21 μM as estimated using a Qubit 4 Fluorometer (Thermo Fisher). The sample was flash-frozen and stored in liquid nitrogen.

For the cryo-EM grid preparation, CHAPSO (3-[(3-cholamidopropyl)dimethylammonio]-2-hydroxy-1-propanesulfonate) was added to the SufR:*P_sufR_* sample at a final concentration of 6 mM immediately before the grid sample preparation. 3.5 μl aliquots of the samples were placed on a 60 second glow-discharged holey carbon grids (Quantifoil Cu R1.2/1.3, 300 mesh). The grids were flash-frozen in liquid ethane using a Vitrobot Mark IV (Thermo Fischer Scientific) under the conditions of 4°C, ≥85% relative humidity and with a blotting time of 4 s. To minimize oxidative damage to the SufR [4Fe-4S] cluster, the Vitrobot chamber was purged with ultrapure nitrogen gas bubbling through a beaker containing Milli-Q water to introduce a mixture of nitrogen gas and water vapor to the chamber during the grid preparation. This strategy preserves relatively high humidity within the chamber while effectively lowering oxygen levels. The oxygen concentration was maintained at ≤4%, as estimated by a Maxtec Handi+ Oxygen Analyzer (Broward A&C Medical).

The cryo-EM data were collected using EPU software on a Thermo Scientific Glacios 2 transmission electron microscope operated at 200 kV. The images were recorded using a Falcon 4i direct detector equipped with a Selectris energy filter, at a nominal magnification of 165,000x, corresponding to a calibrated pixel size of 0.68 Å with a defocus range of -0.8 to -2.5 µm. A total of 13,752 raw movies were collected. The recorded micrographs were saved in the electron-event representation (EER) format and captured at a total dose of 50 electrons/Å² over an exposure time of 4.19 seconds for each micrograph.

### Cryo-EM image data processing

CryoSPARC v4.5 was used for data processing ^54^, following the processing workflow shown in Supplementary Fig. S3. Briefly, after initial processing via motion correction and local CTF correction, 8,496 micrographs were accepted by manual curation. An initial volume of SufR:*P_sufR_* was generated from blob picking using a particle diameter of 150–200 Å and iterative 2D classification. The particles were extracted using a box size of 512 pixels. Template picking and subsequent 2D and 3D iterative refinement yielded a final particle stack of 158,007 particles used to generate the *ab initio* reconstruction (Supplementary Fig. 3). No symmetry was imposed during data processing. The resolution of the final sharpened map was estimated to be 3.5 Å, as measured by gold-standard Fourier shell correlation (GSFSC) (Supplementary Fig. S4). Supplementary Fig. S4 contains additional data regarding map quality. This map was used to build the initial model as described below. A single map was generated from the unmasked half maps using density modification (phenix.resolve_cryo_em), which was used for further refinement of the model and shown in Fig. 3B-C and Supplementary Figs. S5A-B, S6A-B and S10.

### Structural model building and refinement

Real space refinement of the model to the maps was performed in Phenix (v1.21) ^55^. The *Phenix.predict_and_build* tool was used to build the initial model of *Mtb* SufR_6-229_ dimer ^30^, which includes residues 19–228 in one protomer, and residues 18–52 and 83–228 in the second protomer. The model was then manually modified in Coot (v0.9.8), followed by refinement using Phenix.real_space_refine ^56,57^. The *P_sufR_* DNA duplex was built *de novo* in Coot ^58^, manually adjusted and refined in Phenix. The [4Fe–4S] clusters were modeled into the electron density at the C-terminal domain of SufR in Coot and refined in Phenix with geometric restraints for the ligating residues (C179, C192, E195 and C220). The final model was validated using MolProbity (implemented in Phenix) ^59^. The structural statistics are summarized in Supplementary Table 2.

### O_2_ sensitivity test

UV-visible absorption spectroscopy was used to monitor the [4Fe-4S] cluster loss in SufR alone and the DNA-bound SufR. The reducing agent in the purified SufR sample was removed by passing through a desalting column pre-equilibrated with the oxygen-free buffer containing 25 mM Tris, pH 7.5, 150 mM NaCl under anaerobic conditions. The DNA-bound SufR was prepared by mixing SufR with the 50-bp *P_sufR_* DNA in a 1:1.5 molar ratio and the mixture was incubated for 30 minutes under anaerobic conditions. For the O_2_ sensitivity test, the sample containing 200 μM SufR (without and with DNA) was mixed with the air-saturated buffer (25 mM Tris, pH 7.5, 150 mM NaCl) in 1:9 volume ratio, resulting in a final concentration of SufR of 20 μM. The UV–visible absorption spectra were collected at 10-minute intervals for the duration of 1-2 hours as indicated.

### Bioinformatic analysis

For sequence and phylogenetic analyses of representative protein sequences of the COG2345 family, the 400 representative protein sequences were obtained from the Conserved Domain Database ^60^. After removing the sequences from MULTISPECIES, the resulting 326 sequences were aligned using MAFFT (v7.149b) ^61^. The alignment was used to construct a phylogenetic tree using IQ-TREE (v.2.3.4) with the WAG+F+R8 substitution model (determined using MFP) and 100 bootstraps ^62,63^. The resulting tree was visualized and annotated using iTOL^64^.

Phylum classification for the species in the tree was done using the NCBITaxa module in the ETE Toolkit 3.0 ^65^. The sequence logo was generated using WebLogo (v3.7.12) from the aligned 326 representative sequences against *Mtb* SufR (Rv1460) by MAFFT ^61,66^. The gaps in the aligned sequences were removed for clarity.

Representative SufR homologs were selected from those listed in the previous studies and the BioCyc database, as well as blast search using the SufR sequences from *Mtb* and *Sav* as the seeds to obtain diverse SufR homologs in Actinobacteria and Cyanobacteria ^15–17,67^. The sequence logo was then generated as described above.

### 3D structural modeling and alignment

The structural models for the representative COG2345 family proteins were generated using AlphaFold v3.0 ^68^, without and with a [4Fe-4S] cluster (PDB ligand code SF4). The 50-bp *P_sufR_* and the 43-bp *narKGHJI* promoter DNA (*P_narK_*), which includes the protected region in previously reported DNaseI footprinting assay of Cgl ArnR ^42^, was used for generating the structural model of DNA-bound *Mtb* SufR and *Cgl* ArnR, respectively, as described above.

For 3D structural alignment, the top-ranked AlphaFold model for each sequence was used for 3D structural alignment against the DNA-binding domain (DBD) and the sensory domain (SD), respectively, of *Mtb* SufR in the SufR:*P_sufR_* cryo-EM structure using the PyMol Molecular Graphics System v3.0 (https://pymol.org). Of note, the structural models of the DBDs of free *Mtb* SufR and other COG2345 family proteins without DNA show a different conformation of the DBDs relative to the rest of the protein compared with the DNA-bound form, making it impossible to direct 3D structural alignment including the three domains.

### Quantification, statistical analysis and data visualization

The buried surface areas (BSAs) between protein and DNA interface were calculated using the online server PDBePISA ^69^. DNA structural analyses were calculated using Curves+ ^70^. The 3D structural figures were prepared with the PyMol Molecular Graphics System v2.3 (http://www.pymol.org) and ChimeraX ^71^. The interactions between SufR and DNA were visualized using DNAproDB ^72^ with modifications in Adobe Illustrator for figure preparation. The color scheme is shown in the figure legend and briefly described as follows: the functional groups of the nucleotides (N=A, T, G, C) in contact with a protein residue are highlighted: sugars are colored yellow, phosphates in orange. The bases in contact with the protein from the minor grooves are highlighted in pink, and the major grooves in cyan. The rest of the bases are in gray. The base-base stacking is shown by a gray line. All the base pairs in *P_sufR_* are Watson-Crick base pairs (W.C. BP) highlighted by black lines. The protein residues in contact with DNA are highlighted by the colored shapes based on the secondary structure where the residue is located: helices are highlighted by solid red spheres and loops by blue squares. The hydrophilic interactions between a protein residue and the functional group of the nucleotides are indicated by a dashed line, while the hydrophobic and Van der Waals interactions are indicated by a solid line. The color of the lines corresponds to the contacting functional group of the nucleotide. Cavity analysis was performed using the Caver 3.0 software with the probe radii of 1 Å ^73^.

## DATA AVAILABILITY

The cryo-EM maps and the atomic coordinates for the SufR:*P_sufR_* complex have been deposited into the Electron Microscopy Data Bank under accession number EMD-72511 and Protein Data Bank under accession number 9Y5K, respectively. Source data are provided with this paper.

## Supporting information

Validation report

## ACKNOWLEDGEMENTS

The authors thank Dr. Zhi Chen from China Agricultural University for the plasmid used for bioluminescence assay; Dr. Patricia J Kiley from the University of Wisconsin-Madison for the *E. coli BL21(DE3)Suf++ s*train for protein expression; Dr. Huilin Li from Van Andel Institute, Drs. Yihe Huang and Mark Wilson from the University of Nebraska-Lincoln (UNL), and Drs. Jens Kaiser and Xiang Feng at the California Institute of Technology (Caltech) for the insightful discussions; Dr. Songye Chen at the Caltech Cryo-EM Facility and Dr. Eduardo Romero Camacho at the UNL Cryo-EM Facility for their assistance during initial sample screening. We thank the staff at Pacific Northwest Center for Cryo-EM (PNCC) for their assistance during the cryo-EM sample screening and data collection. This work was supported by grants from the National Institutes of Health (R35 GM138157) and the National Science Foundation (Award No. 2033441 and CLP 1846908) to L-M.Z. The content is solely the responsibility of the authors and does not necessarily represent the official views of the National Institutes of Health and the National Science Foundation. The cryo-EM grid preparation and data collection at PNCC was supported by the NIH grant (R24GM154185). This work was completed utilizing the Holland Computing Center (HCC) of the University of Nebraska, which receives support from the UNL Office of Research and Innovation and the Nebraska Research Initiative.

## AUTHOR CONTRIBUTIONS

L-M.Z. directed the project. L-M.Z., Z. L. and T.W. contributed to experimental design. Z.L., L.Z. and T.W. conducted molecular biology work, expressed, purified and characterized the proteins. O.D., M.D.F. and Z.L. conducted the cryo-EM grid preparation and data collection. C.O. and L-M.Z. carried out the bioinformatic analysis and structural modeling. Z.L. and L-M.Z conducted the cryo-EM data analysis. Z.L., C.O. and L-M.Z. wrote the manuscript. All authors contributed to the manuscript preparation.

