## Supplementary Materials for "A New Paradigm of Transcriptional Regulation by the SufR-Like Iron-Sulfur Transcription Factors"

### Supplementary Tables

**Supplementary Table 1.** Bacterial strains, plasmids and oligos used in this study.

| **Label** | **Description** | **Reference** |
|---|---|---|
| **Strains** | | |
| ***E. coli*** | | |
| XL1-Blue | Host strain for general clone | Stratagene |
| BL21(DE3) Suf++ | Host strain for recombinant protein expression, received from Prof. Patricia J Kiley (University of Wisconsin-Madison). | [1] |
| JM109(DE3) | Host strain for bioluminescence test | Promega |
| **Plasmids** | | |
| **Plasmids for expression and purification of protein from *E. coli*** | | |
| pET28a - MtbSuf $R_{6-229}$- $His_6$ | A DNA fragment encoding residues 6-229 ($SufR_{6-229}$) was amplified from *Mycobacterium tuberculosis* (*Mtb*) H37Rv genomic DNA (locus no: Rv1460) and inserted into the NcoI/Xho sites of pET28a(+) for expressing *Mtb* $SufR_{6-229}$ -$His_6$ used for the cryo-EM structural analysis, *in vitro* assays (EMSA, MST and SEC/MALS) and the heterologous lux-reporter system; $Kan^R$ | This study |
| pET28a-MtbSufR-M1 | A modification of pET28a-Mtb$SufR_{6-229}$-$His_6$ by site-directed mutagenesis for expressing $SufR_{6-229}$-$His_6$ with the triple mutation of R50A, R51A and H52A used for EMSAs and the heterologous lux-reporter system; $Kan^R$ | This study |
| pET28a-MtbSufR-M2 | A modification of pET28a-Mtb$SufR_{6-229}$-$His_6$ by site-directed mutagenesis for expressing $SufR_{6-229}$-$His_6$ with the double mutation of R75A and R77A used for EMSAs and the heterologous lux-reporter system; $Kan^R$ | This study |
| pET28a-MtbSufR-R51A | A modification of pET28a-Mtb$SufR_{6-229}$-$His_6$ by site-directed mutagenesis for expressing $SufR_{6-229}$-$His_6$ with a R51A mutation used for EMSAs and the heterologous lux-reporter system; $Kan^R$ | This study |
| pET28a-MtbSufR-E195D | A modification of pET28a-Mtb$SufR_{6-229}$-$His_6$ by site-directed mutagenesis for expressing $SufR_{6-229}$-$His_6$ with a E195D mutation and used for the EMSAs and the heterologous lux-reporter system; $Kan^R$ | This study |
| pET28a-MtbSufR-R75M | A modification of pET28a-Mtb$SufR_{6-229}$-$His_6$ by site-directed mutagenesis for expressing $SufR_{6-229}$-$His_6$ with a R75M mutation used for the EMSAs and the heterologous lux-reporter system; $Kan^R$ | This study |
| **Plasmids for bioluminescence test** | | |
| pOsufRlux | The pOsufRlux plasmid received from Prof. Zhi Chen (China Agricultural University) was used for the heterologous lux-reporter system with modification; $Kan^R$ | [2] |
| pCS-*PsufR*-Lux-Amp | This plasmid was modified from pOsufRlux. The $Kan^R$ marker in pOsufRlux was replaced by an $Amp^R$ marker amplified from pET21b. The promoter of the *suf* operon from *Mtb* H37Rv genomic DNA (genomic region 1,645,886-1,646,133, GenBank accession number: NC_000962.3) was inserted into the BamHI/XhoI site of the pOsufRlux. This plasmid was used together with pET28a-Mtb$SufR_{6-229}$-$His_6$ (wildtype or mutants) in the heterologous lux-reporter system; $Amp^R$ | This study |
| pCS-*PsufR*-M2-Lux-Amp | A modification of pCS- *PsufR*-Lux-Amp by mutating the 3'-end AT-rich region of the promoter (see the oligo sequence $P_{sufR\text{-}M2}$). This plasmid was used together with the pET28a-Mtb$SufR_{6-229}$-$His_6$ in the heterologous lux-reporter system; $Amp^R$ | This study |

| | | |
|---|---|---|
| pCS-*PsufR*-DM-Lux-Amp | A modification of pCS- *PsufR*-Lux-Amp by mutating both AT-rich regions of the promoter (see the oligo sequence $P_{sufR\text{-}DM}$). This plasmid was used together with the pET28a-MtbSufR$_{6\text{-}229}$-His$_6$ in the heterologous lux-reporter system; Amp$^R$ | This study |
| pCS- *PsufR*-PM-Lux-Amp | A modification of pCS- *PsufR*-Lux-Amp by mutating the palindromic sequence of the promoter (see the oligo sequence $P_{sufR\text{-}PM}$). This plasmid was used together with the pET28a-MtbSufR$_{6\text{-}229}$-His$_6$ in the heterologous lux-reporter system; Amp$^R$ | This study |
| **Oligos used for the cryo-EM analysis and DNA-binding assays for SufR** (the palindromic sequences are highlighted in blue, and the mutated sites are in red) | | |
| $P_{sufR}$ | Forward:<br>5'-<br>CCGATCAACGGAATTTTGTCACACTGATGTTGTGAAAATCCCGGCGG<br>TCT-3',<br>5'-Cy5 labeled for the MST and EMSAs<br><br>Reverse: 5'-<br>AGACCGCCGGGATTTTCACAACATCAGTGTGACAAAATTCCGTTGAT<br>CGG-3'<br>Used for EMSAs, MST and cryo-EM structural analysis | This study |
| $P_{sufR\text{-}M1}$ | Forward:<br>5'-<br>CCGATCAACGGAACTGTGTCACACTGATGTTGTGAAAATCCCGGCGG<br>TCT-3'<br>5'-Cy5 labeled for the EMSAs<br>Reverse: 5'-<br>AGACCGCCGGGATTTTCACAACATCAGTGTGACACAGTTCCGTTGAT<br>CGG-3'<br><br>Used for EMSAs | This study |
| $P_{sufR\text{-}M2}$ | Forward:<br>5'-<br>CCGATCAACGGAATTTTGTCACACTGATGTTGTGACAGTCCCGGCGG<br>TCT-3'<br>5'-Cy5 labeled for EMSAs<br><br>Reverse: 5'-<br>AGACCGCCGGGACTGTCACAACATCAGTGTGACAAAATTCCGTTGAT<br>CGG-3'<br><br>Used for EMSAs | This study |
| $P_{sufR\text{-}DM}$ | Forward:<br>5'-<br>CCGATCAACGGAACTGTGTCACACTGATGTTGTGACAGTCCCGGCGG<br>TCT-3'<br>5'-Cy5 labeled for EMSAs<br><br>Reverse: 5'-<br>AGACCGCCGGGACTGTCACAACATCAGTGTGACACAGTTCCGTTGAT<br>CGG-3'<br><br>Used for EMSAs | This study |

| | | |
|---|---|---|
| $P_{sufR\text{-}PM}$ | Forward:<br>5'-<br>CCGATCAACGGAATTTTGTCGAAGTGATGTCACTAAAATCCCGGCGG<br>TCT-3'<br>5'-Cy5 labeled for EMSAs<br>Reverse:<br>5'-<br>AGACCGCCGGGATTTTAGTGACATCACTTCGACAAAATTCCGTTGAT<br>CGG-3'<br>Used for EMSAs | This study |
| $P_{sufR\text{-}31bp}$ | Forward:<br>5'-AATTTTGTCACACTGATGTTGTGAAAATCCC-3',<br>Reverse:<br>5'-GGGATTTTCACAACATCAGTGTGACAAAATT-3'<br><br>5'-Cy5 labeled for the EMSAs | This study |
| **Oligos used for the DNA-binding assays for *Cgl* ArnR**<br>(the consensus sequences are highlighted in bold fonts, the protected sequences in the DNaseI footprinting assays are colored blue, and the mutated sites are in red) | | |
| $P_{narK}$ | Forward:<br>5'-GTTGCC**TAATTAAATA**CGGAAAACCCCGTTGAAAAACATGCGG<br>-3',<br>Reverse:<br>5'- CCGCATGTTTTTCAACGGGGTTTTCCG**TATTTAATTA**GGCAAC-3'<br>Used for EMSAs analysis | This study |
| $P_{narK\text{-}M1}$ | Forward:<br>5'-GTTGCC**TAATTGCATA**CGGAAAACCCCGTTGAAAAACATGCGG<br>-3',<br>Reverse:<br>5'- CCGCATGTTTTTCAACGGGGTTTTCCG**TATGCAATTA**GGCAAC-3'<br>Used for EMSAs analysis | This study |
| $P_{narK\text{-}M2}$ | Forward:<br>5'-GTTGCC**TAATTAAATA**CGGAAAACCCCGTTGAACGACATGCGG<br>-3',<br>Reverse:<br>5'- CCGCATGTCGTTCAACGGGGTTTTCCG**TATTTAATTA**GGCAAC-3'<br>Used for EMSAs analysis | This study |
| $P_{narK\text{-}DM}$ | Forward:<br>5'-GTTGCC**TAATTGCATA**CGGAAAACCCCGTTGAACGACATGCGG<br>-3',<br>Reverse:<br>5'- CCGCATGTCGTTCAACGGGGTTTTCCG**TATGCAATTA**GGCAAC-3'<br>Used for EMSAs analysis | This study |

**Supplementary Table 2.** Cryo-EM data collection, refinement and validation statistics.

| | |
|---|---|
| **Data Collection and processing** | |
| Microscope | Glacios 2 |
| Voltage (kV) | 200 |
| Detector | Falcon 4i |
| Magnification | 165,000 |
| Pixel size (Å) | 0.68 (physical resolution) |
| Electron dose ($e^-$/Å$^2$) | 50 |
| Exposure time (s) | 4.19 |
| Defocus range (μm) | -0.8 ~ -2.5 |
| Number of micrographs | 13752 |
| Software | CryoSPARC 4.5 |
| Final micrographs used | 13752 |
| Initial particles | 5,769,627 |
| Final particles | 172,193 |
| Symmetry imposed | C1 |
| Map resolution (Å) | 3.48 Å |
| FSC threshold | 0.143 |
| Map sharpening B-factor (Å$^2$) | 136.5 |
| **Model Refinement** | |
| Refinement software | Phenix and coot |
| Initial model | de novo |
| Model-to-map resolution (Å) | 3.0/3.2/3.48 |
| FSC threshold | 0.143/0.5/0.85 |
| **Model Composition** | |
| Non-hydrogen atoms | 4673 |
| Protein residues | 424 (chain A: 17-228, chain B: 17-228) |
| Nucleotide | 74 (chain C: 1-37, chain D:1-37) |
| Ligands | 2 (SF4) |
| **RSMDs** | |
| Bond lengths (Å) | 0.004 |
| Bond angles (°) | 0.555 |
| **B factors (Å$^2$) (Min/Max/Mean)** | |
| Protein residues | 10.03/150.54/52.14 |
| Nucleotides | 46.75/191.47/108.60 |
| Ligands | 28.03/79.60/48.85 |
| **Validation** | |
| MolProbity score | 1.83 |
| Clashscore | 5.57 |
| Rotamer outliers | 1.29 |
| **Ramachandran Plot** | |
| favored (%) | 93.10 |
| allowed (%) | 6.43 |
| outliers (%) | 0.48 |

FSC, Fourier shell correlation; RMSD, root-mean-square deviation.

**Supplementary Table 3.** Summary of the conservation of the sequence motifs for DNA binding and [4Fe-4S] cluster binding among the representative COG2345 members. Related to Fig. 5A.

| DNA binding residues in SufR | | | | | | [4Fe-4S] cluster binding residues in SufR | | | |
|---|---|---|---|---|---|---|---|---|---|
| **R50** | **R51** | **H52** | **G76** | **R77** | **P78** | **C179** | **C192** | **C220** | **E195** |
| R: 83.4% | R: 38.04% | H: 84.66% | G: 99.69% | R: 97.55% | P: 99.08% | C: 99.39% | C: 97.85% | C: 98.77% | E: 63.80% |
| K: 3.07% | Q: 18.10% | Q: 9.20% | R: 0.31% | K: 1.84% | T: 0.61% | A: 0.31% | V: 0.61% | L: 0.31% | H: 19.02% |
| H: 2.15% | K: 7.36% | R: 3.37% | | A: 0.61% | R: 0.31% | N: 0.31% | F: 0.31% | R: 0.31% | N: 7.36% |
| Q: 1.53% | E: 7.05% | T: 0.61% | | | | | L: 0.31% | T: 0.31% | D: 4.60% |
| N: 0.92% | H: 2.76% | S: 0.31% | | | | | R: 0.31% | V: 0.31% | R: 0.92% |
| T: 0.92% | D: 0.92% | Y: 0.31% | | | | | S: 0.31% | | Q: 0.61% |
| S: 0.61% | N: 0.92% | Other non-Polar: 1.53% | | | | | Y: 0.31% | | S: 0.31% |
| Y: 0.61% | T: 0.92% | | | | | | | | Other Nonpolar: 3.37% |
| Other Nonpolar: 6.75% | S: 0.31% | | | | | | | | |
| | Y: 0.31% | | | | | | | | |
| | Other Nonpolar: 23.31% | | | | | | | | |

### Supplementary Figures

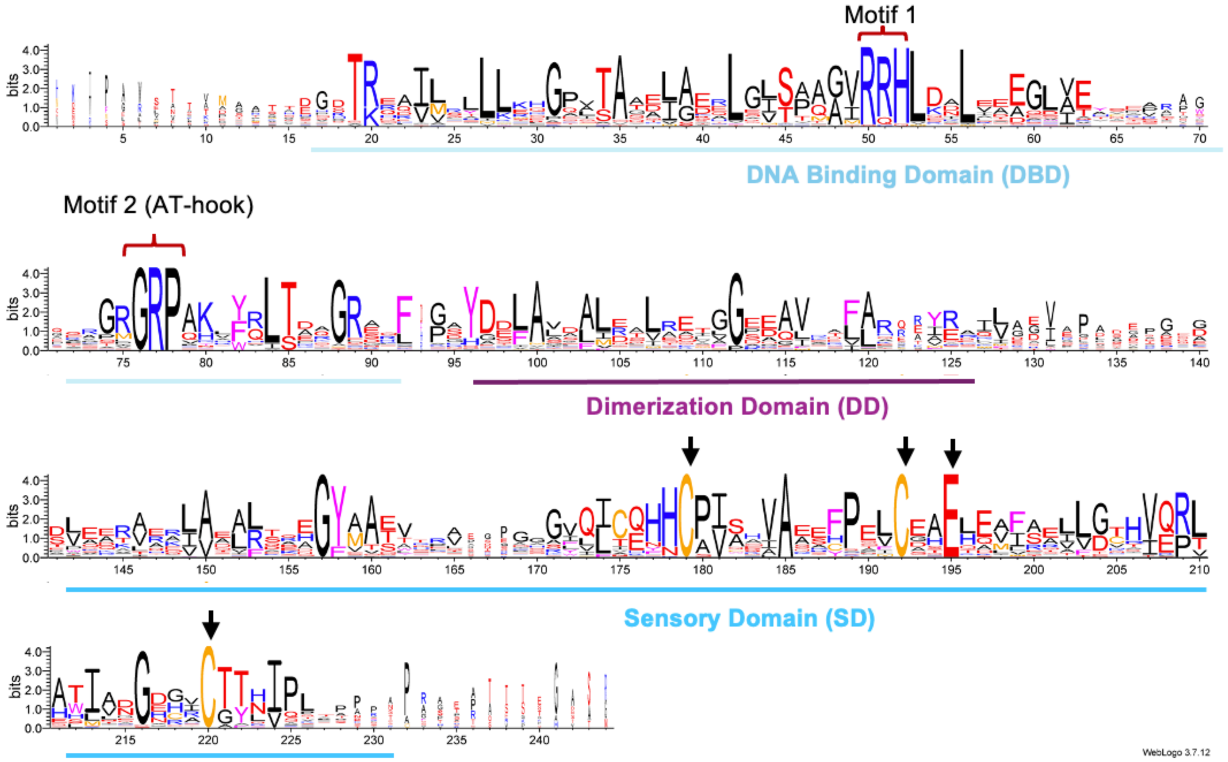


**Supplementary Figure S1.** Sequence logo of selected SufR homologs from Actinobacteria and Cyanobacteria listed in Supplementary DataSet 1 (see in the Methods). The numbering of the residues in the sequence logo follows the *Mtb* Wbl proteins, with alignment gaps in the sequence logos removed for clarity. The two conserved Arg-rich motifs (the "50-RRH-52" motif [Motif1] and the central "76-GRP-78" AT-hook [Motif2]) are highlighted by red brackets. The conserved residues involved in [4Fe-4S] cluster binding in *Mtb* SufR are indicated by black arrows.

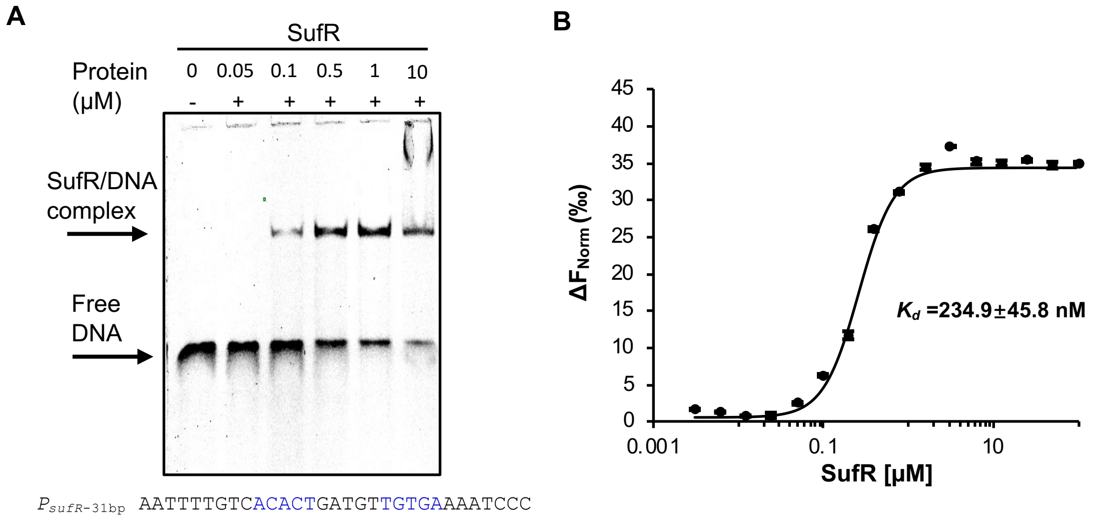


**Supplementary Figure S2. Characterization of SufR binding to the promoter DNA.** (A) Electrophoretic mobility shift assays (EMSAs) of SufR binding to the 31-bp promoter DNA ($P_{sufR\text{-}31bp}$), which was used in the previous EMSAs [3]. (B) Microscale thermophoresis (MST) assay of SufR binding to the 50-bp $P_{sufR}$ DNA. The fitted curve is shown as a solid line, with each data point representing the mean and standard deviation of three independent measurements.

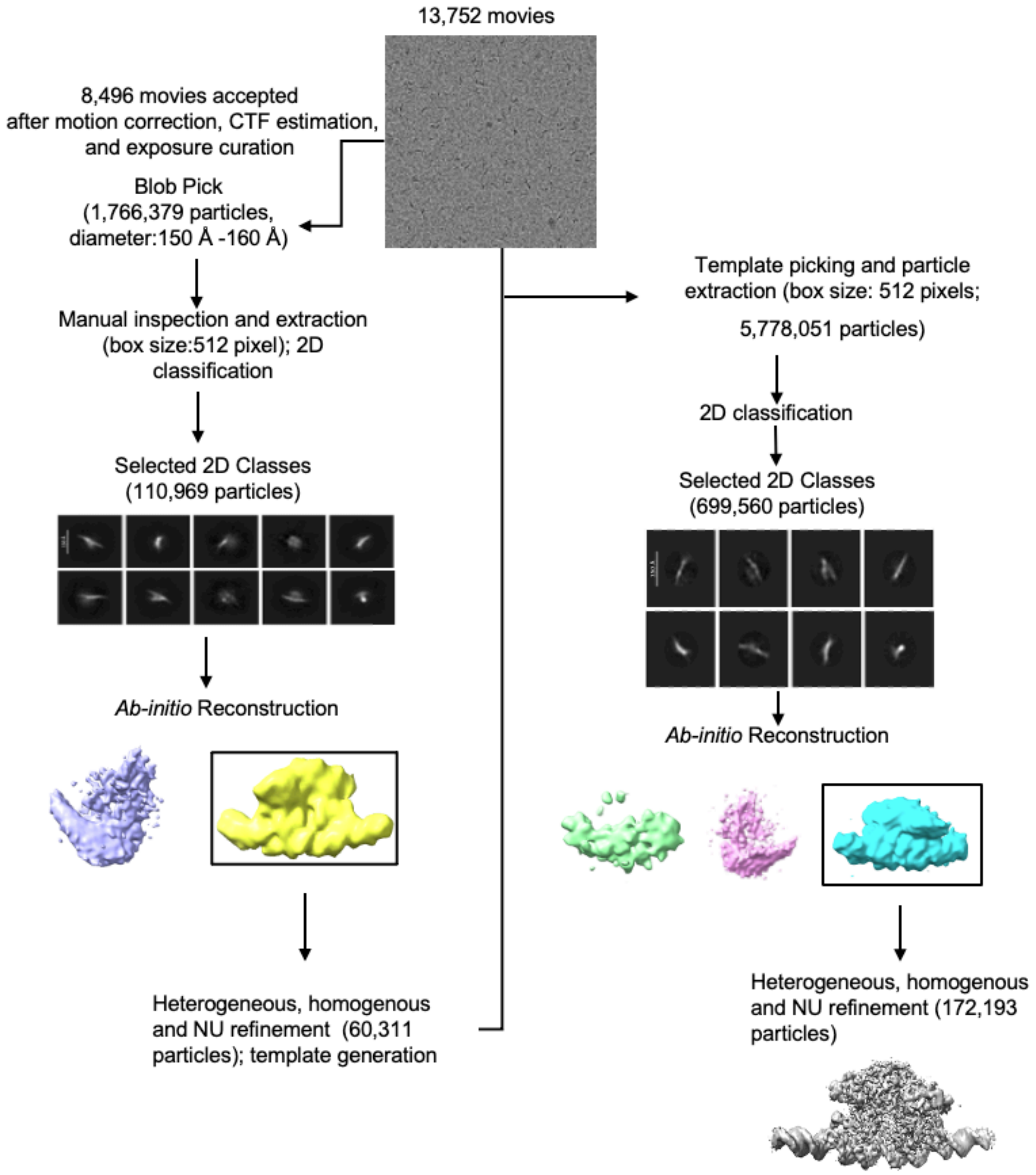


**Supplementary Figure S3. Cryo-EM processing pipeline of the SufR:*$P_{sufR}$* complex.** See the detailed description in the Methods.

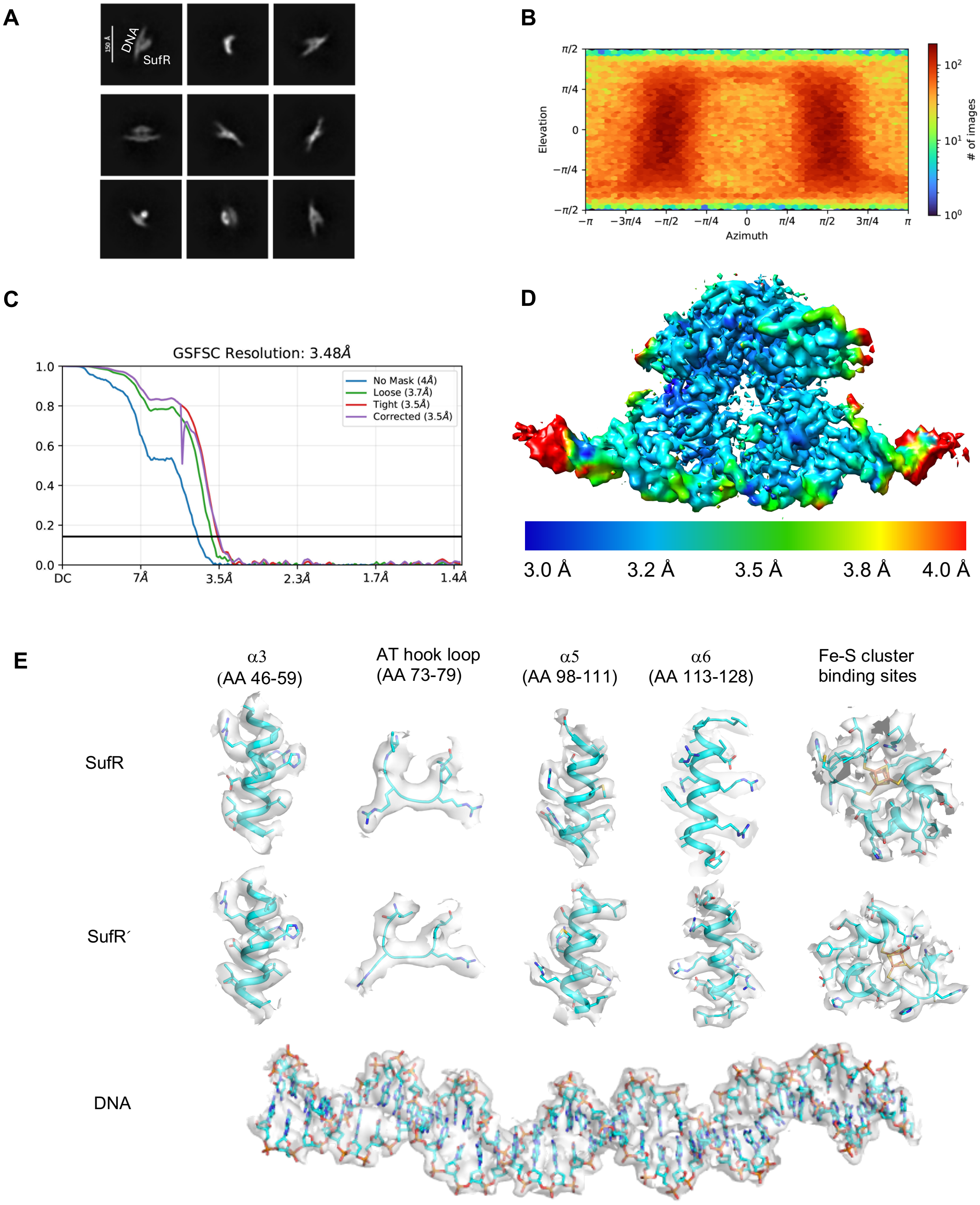


**Supplementary Figure S4. Evaluation of the final cryo-EM density map of the SufR:$P_{sufR}$ complex.** (A) Selected reference-free 2D class averages used in the final 3D reconstruction of the EM map. (B) Angular distribution calculated in cryoSPARC for particles used in the final 3D reconstruction of the EM map. (C) Gold-standard Fourier shell correlation indicates an overall resolution of 3.5 Å. (D) Color-coded local resolution map of the final 3D map. (E) The key structural elements (with the range of the amino acids [AA] shown in the bracket) of the SufR:DNA complex model are fitted within the sharpened cryo-EM map.

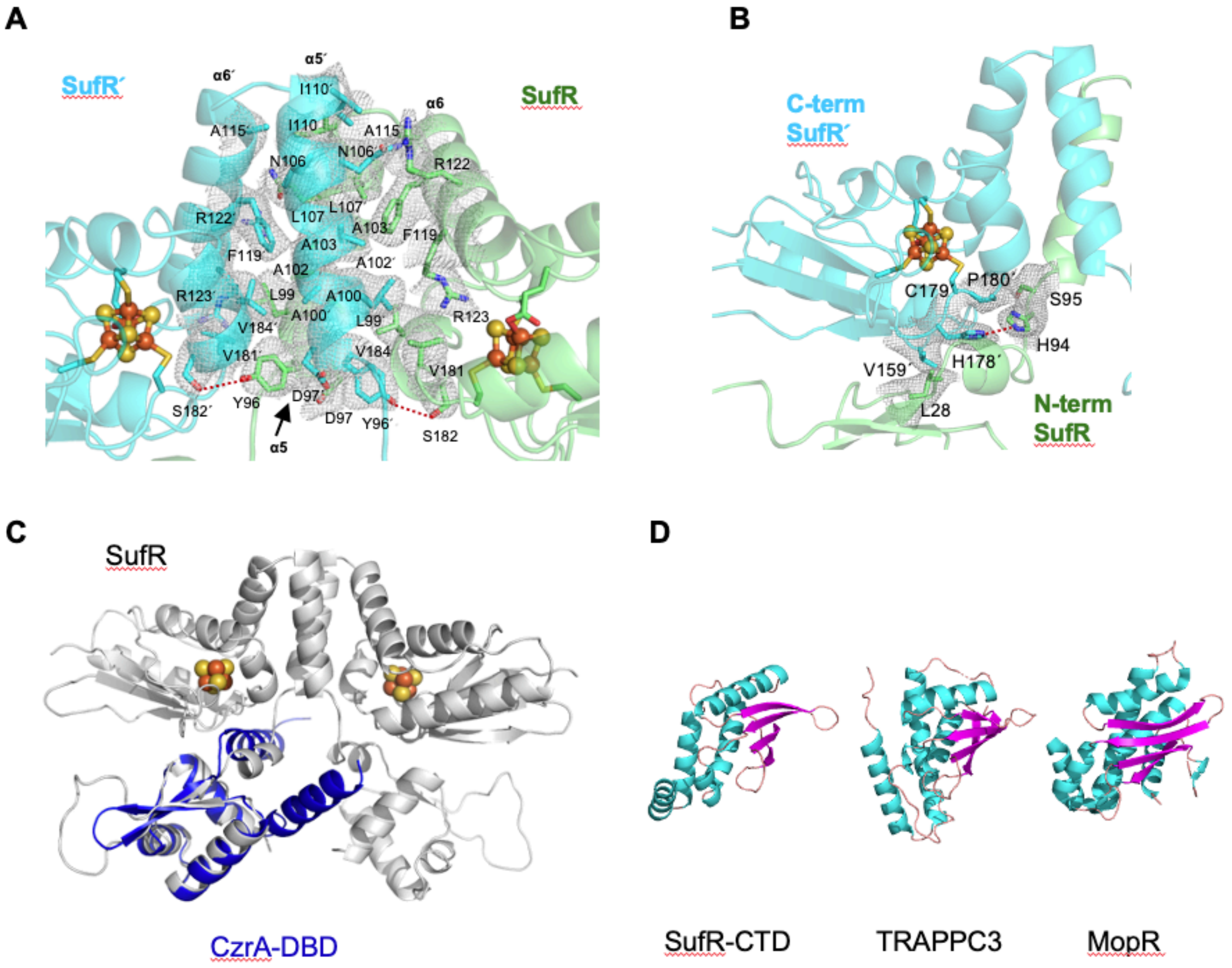


**Supplementary Figure S5. Domain architecture of the SufR:$P_{sufR}$ cryo-EM structure.** Panels (A) and (B) are the highlights of the key residues at the interaction interface between the two dimerization domains and the N-terminus and the domain-swapped C´-terminus of SufR, respectively, with the local cryo-EM density map. The maps are colored gray and contoured at 2 σ. (C) Structural alignment of the DNA-binding domain of the ArsR-family metal-sensing transcriptional repressor CzrA from *Staphylococcus aureus* (PDB ID: 1R1U, blue) against that of SufR (gray), with the $RMSD_{C\alpha}$ of 2.1 Å over 39 residues. The rest of the CzrA structure does not align with SufR and is not shown for clarity. (D) Comparison of the C-terminal domain (CTD, including the dimerization domain and the sensory domain) of SufR with the regulatory subunit of the trafficking protein particle complex subunit 3 (TRAPPC3) (PDB ID:1WC8) and the NtrC family transcription factor MopR (PDB ID: 5KBG). For all structures, the helices are colored in cyan, the ß strands are in magenta and the loops are in brown.

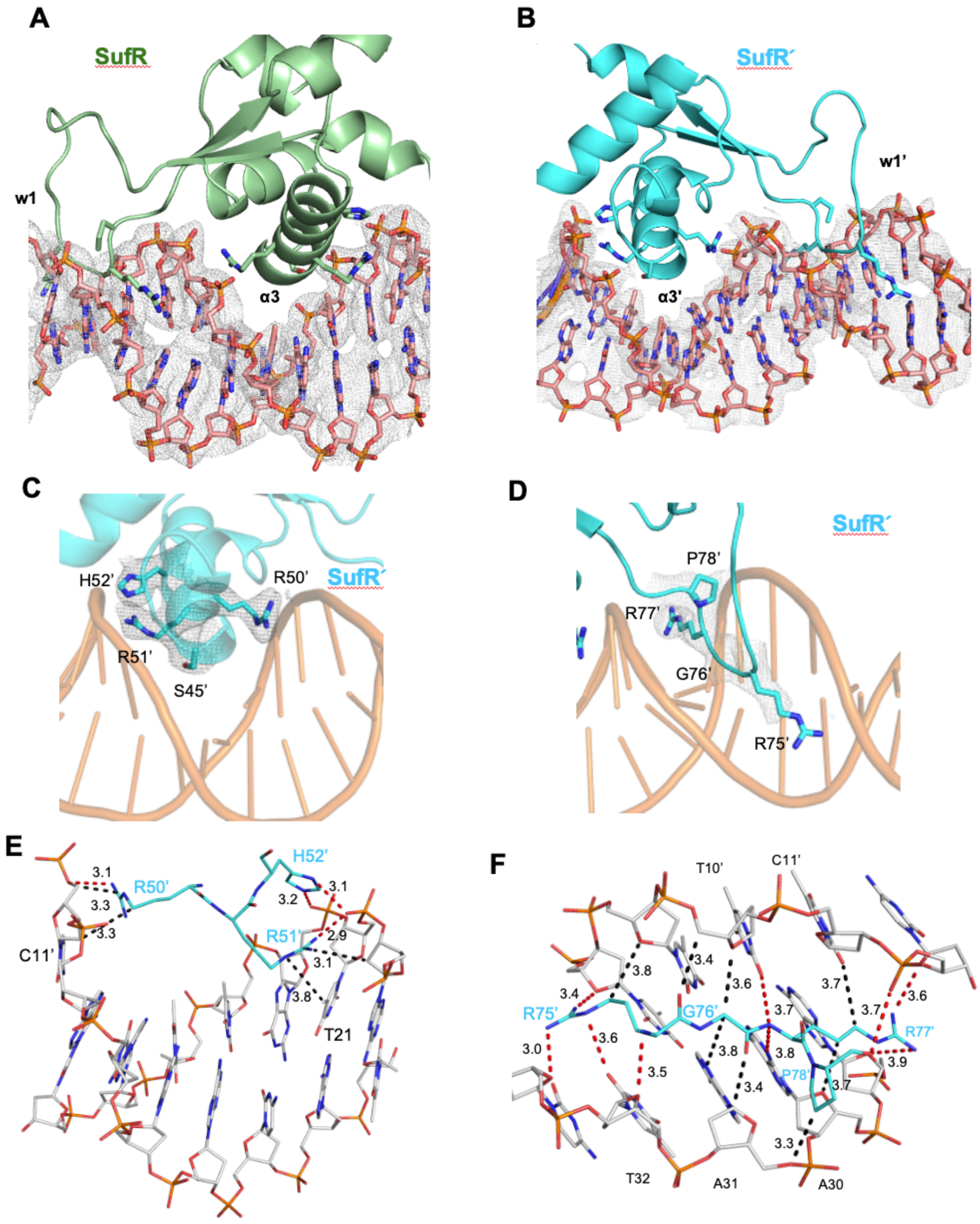


**Supplementary Figure S6. The local cryo-EM density maps and contacts at the SufR:*$P_{sufR}$* interface.** Panels (A) and (B) show the local cryo-EM density map of the DNA fragments around the two SufR binding sites. Panels (C) and (D) show the local cryo-EM density map around the DNA-binding residues in helix α3 and the AT-hook, respectively, of SufR′, compared to Fig. 3B-C. The maps are colored gray and contoured at 2 σ. Panels (E) and (F) are comparisons of the DNA contacts by the "RRH" motif and the "RGRP" motif, respectively, in SufR′. The contacting residues, nucleotide bases and the distances (Å) are marked. The polar interactions are highlighted by red dashed lines, and the non-polar interactions are in black dashed lines.

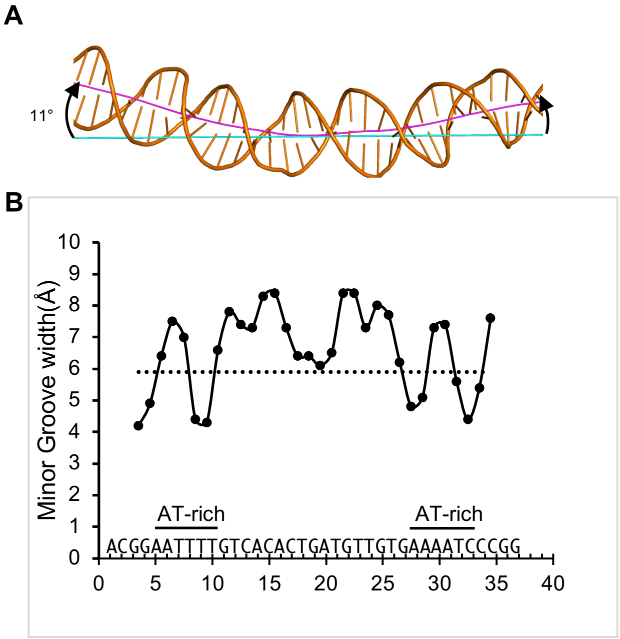


**Supplementary Figure S7. DNA structure analysis of the SufR-bound *$P_{sufR}$*.** (A) The bent SufR-bound $P_{sufR}$ DNA axis (colored magenta) calculated by Curves+, when compared to the straight axis (cyan) illustrating the unbent B-form DNA. (B) Minor-groove widths of the SufR-bound $P_{sufR}$ DNA. The groove width is defined as the distance between the closest phosphates, subtracted by 5.8 Å (the sum of the van der Waals radii of the two phosphate atoms). The dashed line indicates the mean value of canonical minor groove widths in B-form DNA.

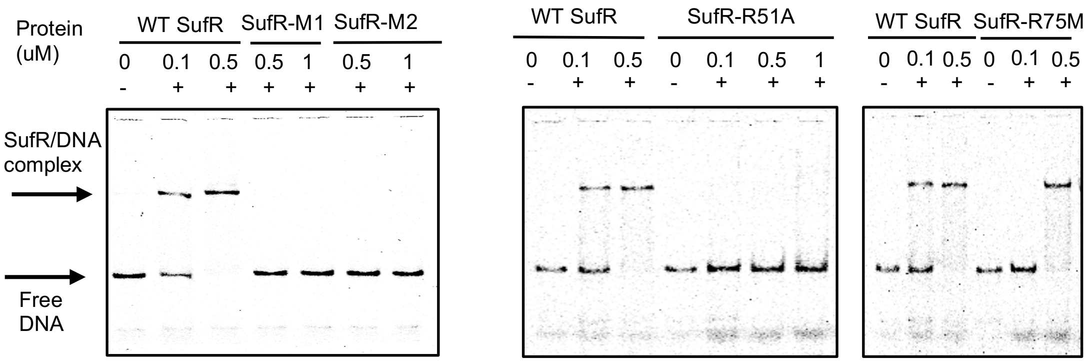


**Supplementary Figure S8**. The uncropped gel images of the EMSAs shown in Figure 3E.

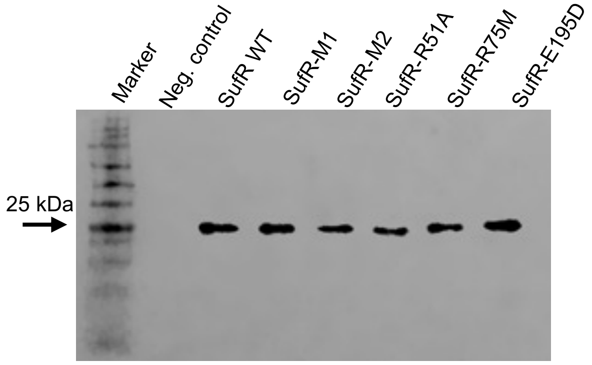


**Supplementary Figure S9**. **Western blot analysis of the *Mtb* SufR$_{6\text{-}229}$-His$_6$ protein (wildtype and mutants as indicated) in *E. coli* JM109(DE3) strains used for the bioluminescence assays.** A sample from the cells with the empty vector was used as the negative control (Neg. control). The theoretical molecular weight of *Mtb* SufR$_{6\text{-}229}$-His$_6$ is 25.1 kDa in the monomeric state.

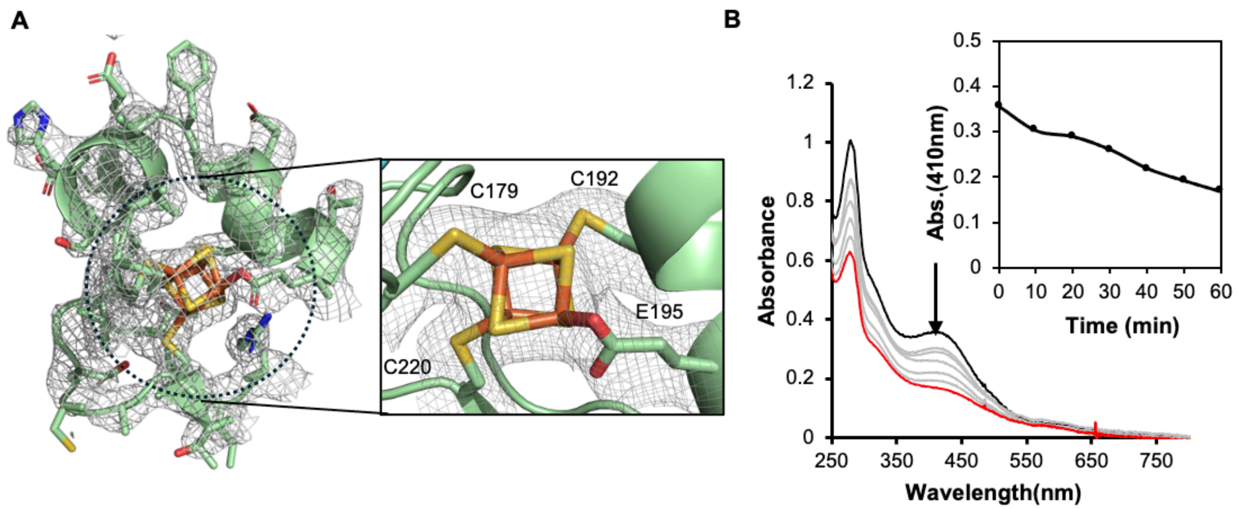


**Supplementary Figure S10. The local environment at the [4Fe-4S] cluster binding pocket of the *P<sub></sub>*$_{sufR}$-bound SufR.** (A) The local cryo-EM maps around the [4Fe-4S] cluster binding pocket, with a zoomed-in view of the first coordination of the cluster binding site. The map is colored gray and contoured at 2 σ. (B) UV-visible absorption spectra of free SufR upon exposure to $O_2$ at 0 min (black curve), between 10-50 min (gray curves) and at 60 min (red curve). Inset, the plot of the absorption intensity at 410 nm as a function of time, indicating loss of the Fe-S cluster.

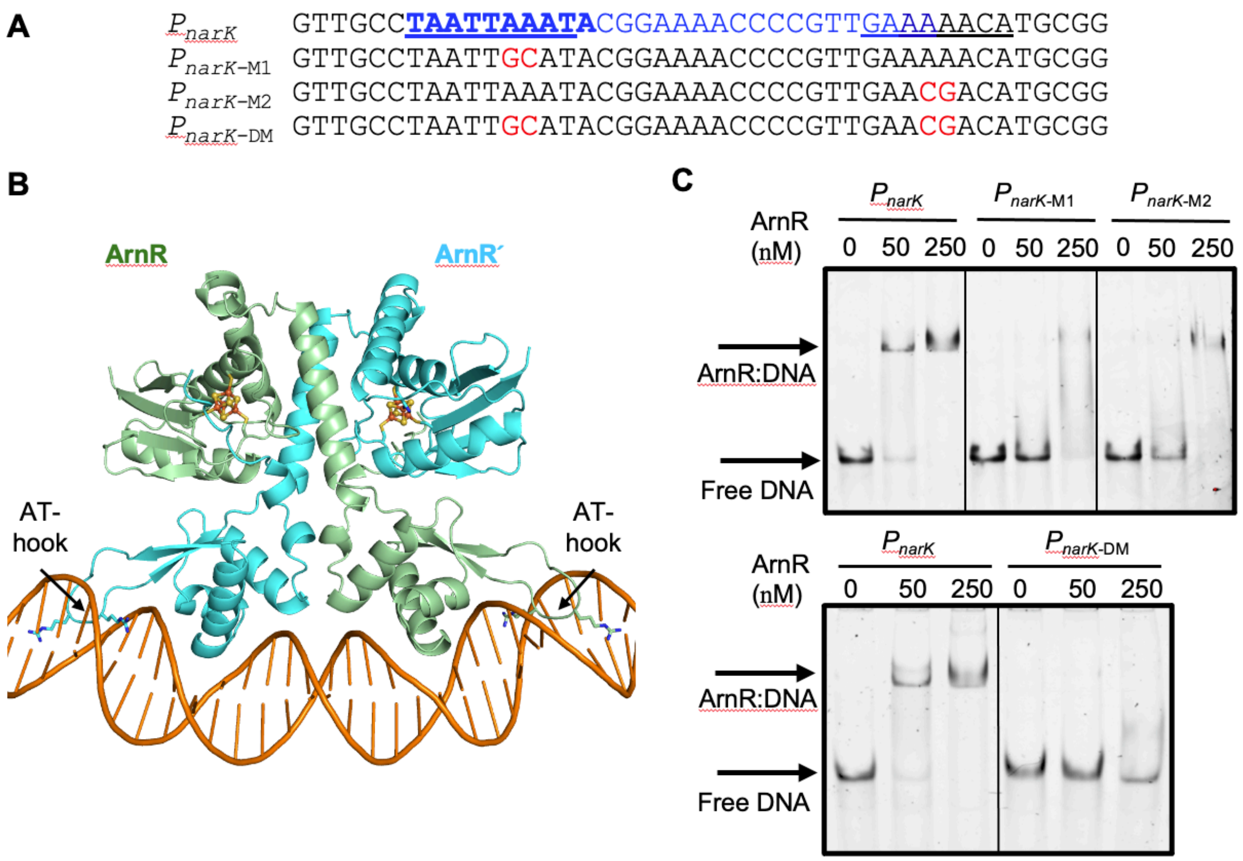


**Supplementary Figure S11. Characterization of *Cgl* ArnR binding to the $P_{narK}$ promoter** (A) The 43-bp $P_{narK}$ sequence used in the AlphaFold modeling (Panel B) and the mutation sequences in the two A/T rich regions used in the electrophoretic mobility shift assays (EMSAs) (Panel C). The protected region in the reported DNaseI footprinting assays[4] is highlighted in blue in $P_{narK}$, with the consensus ArnR-binding sequence (TAWTTAAWTA where W=A/T is shown in bold. The underlined sequences are predicted in contact with the ArnR AT-hooks. The mutated nucleotides are highlighted in red. (B) Overview of the AlphaFold model of ArnR bound to $P_{narK}$. The [4Fe-4S] clusters are highlighted in ball and sticks, and the AT-hook residues at the protein:DNA interface are shown in sticks. The Fe atoms are colored red, S in orange, N in blue and O in red. The TM score of the AlphaFold model is 0.84. (C) EMSAs of ArnR binding to the $P_{narK}$ (wildtype and mutant sequences as indicated and shown in Panel A). 50 nM of the unlabeled DNA was used in the assays (see in the Materials and Methods).

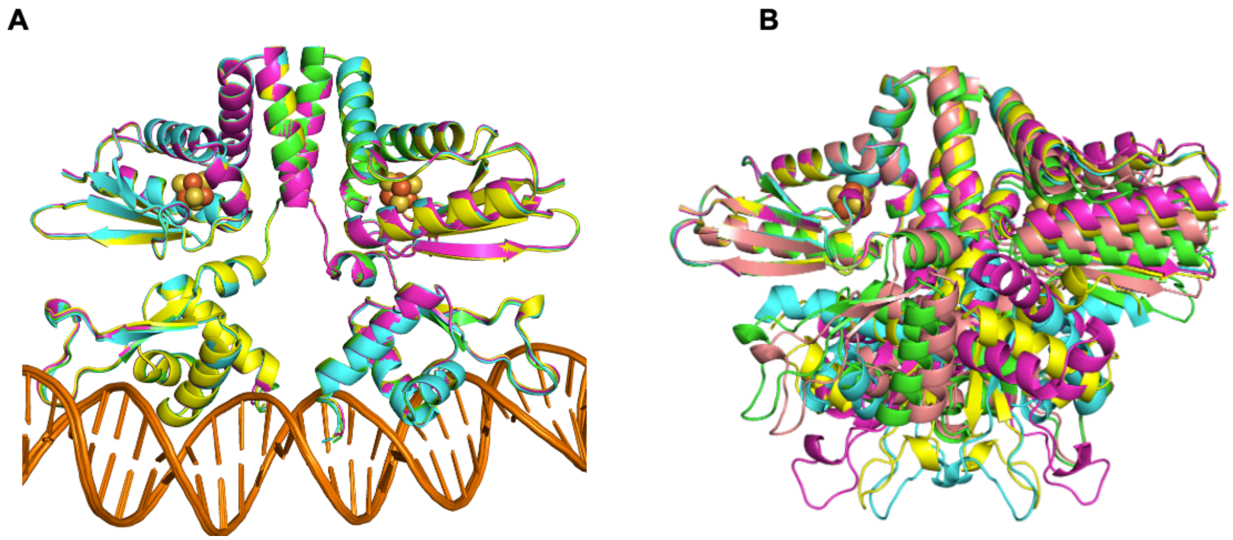


**Supplementary Figure S12. Comparison of the AlphaFold models of *Mtb* SufR.** (A) Comparison of the five models of *Mtb* SufR bound to the $P_{sufR}$ promoter. The TM score of the AlphaFold model is 0.76. (B) Comparison of the five models of free *Mtb* SufR without DNA. The TM score of the AlphaFold model is 0.49. In all the models, DNAs are colored orange, while the five SufR models are colored orange, magenta, yellow, cyan and green, respectively.

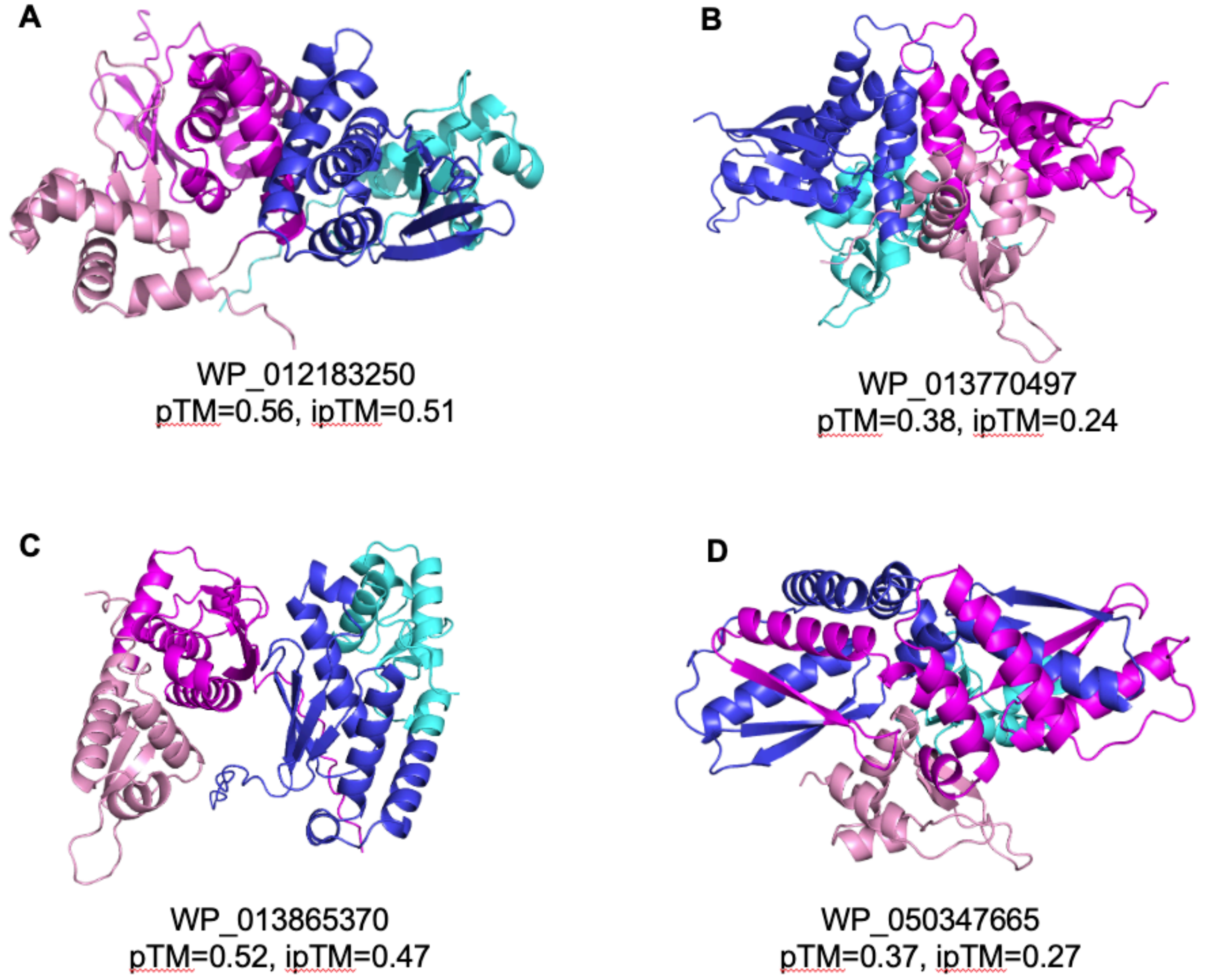


**Supplementary Figure S13. AlphaFold models of four COG2345 outliers in the domain architecture, using the overall architecture of the dimerization domain (DD) and the sensing domain in SufR as the reference.** In all the cases, only the top-ranked model is shown for clarity. The DBDs are colored pink and cyan, respectively, while the rest of the domains are colored magenta and blue, respectively. The low confidence scores in the accuracy of the predicted relative positions of the subunits within the complex (ipTM <0.6) indicate the high uncertainties of these *ab initio* models.
